# CSF1R-dependent macrophages control postnatal somatic growth and organ maturation

**DOI:** 10.1101/2020.11.29.402859

**Authors:** Sahar Keshvari, Melanie Caruso, Ngari Teakle, Lena Batoon, Anuj Sehgal, Omkar L. Patkar, Michelle Ferrari-Cestari, Cameron E. Snell, Chen Chen, Alex Stevenson, Felicity M. Davis, Stephen J. Bush, Clare Pridans, Kim M. Summers, Allison R. Pettit, Katharine M. Irvine, David A. Hume

## Abstract

Homozygous mutation of the *Csf1r* locus (*Csf1rko*) in mice, rats and humans leads to multiple postnatal developmental abnormalities. To enable analysis of the mechanisms underlying the phenotypic impacts of *Csf1r* mutation, we bred a rat *Csf1rko* allele to the inbred dark agouti (DA) genetic background and to a *Csf1r*-mApple reporter transgene. The *Csf1rko* led to almost complete loss of embryonic macrophages and ablation of most adult tissue macrophage populations. We extended previous analysis of the *Csf1rko* phenotype to early postnatal development to reveal impacts on musculoskeletal development and proliferation and morphogenesis in multiple organs. Expression profiling of 3-week old wild-type (WT) and *Csf1rko* livers identified 2760 differentially expressed genes associated with the loss of macrophages, severe hypoplasia, delayed hepatocyte maturation, disrupted lipid metabolism and the IGF1/IGF binding protein system. Older *Csf1rko* rats developed severe hepatic steatosis. Consistent with the developmental delay in the liver *Csf1rko* rats had greatly-reduced circulating IGF1. Transfer of WT bone marrow (BM) cells at weaning without conditioning repopulated resident macrophages in all organs, including microglia in the brain and reversed the mutant phenotypes enabling long term survival and fertility. WT BM transfer restored osteoclasts, eliminated osteopetrosis, restored bone marrow cellularity and architecture and reversed granulocytosis and B cell deficiency. *Csf1rko* rats had an elevated circulating CSF1 concentration which was rapidly reduced to WT levels following BMT. However, CD43^hi^ non-classical monocytes, absent in the *Csf1rko*, were not rescued and bone marrow progenitors remained unresponsive to CSF1. The results demonstrate that the *Csf1rko* phenotype is autonomous to BM-derived cells and indicate that BM contains a progenitor of tissue macrophages distinct from hematopoietic stem cells. The model provides a unique system in which to define the pathways of development of resident tissue macrophages and their local and systemic roles in growth and organ maturation.

## Introduction

Resident macrophages are abundant in every tissue in the body and adapt to each tissue environment by expressing unique gene sets required for their local functions (reviewed in [1, 2]). Their differentiation from progenitor cells, their gene expression profile and their survival/maintenance in tissues is controlled by two ligands, colony stimulating factor 1 (CSF1) and interleukin 34 (IL34), which each signal through the CSF1 receptor (CSF1R) [3]. The biology of CSF1R and its ligands is conserved from birds to mammals [4–6]. In mice, the impacts of a homozygous *Csf1r* knockout mutation (*Csf1rko*) include perinatal mortality, postnatal growth retardation, increased bone density (osteopetrosis), global defects in brain development and abnormalities of the sensory nervous system, infertility and delayed pancreatic beta cell development (reviewed in [3, 7]). Many of the effects of the *Csf1rko* in mice are shared with mutations in the *Csf1* gene [3, 7].

Phenotypes associated with biallelic recessive mutations in human *CSF1R* appear somewhat less severe although it is not clear that any such mutations are definitively null for CSF1R function [6]. Patients present with abnormal skeletal development and calcification, ventricular enlargement (hydrocephalus) and selective loss of microglia in the brain leading to degenerative encephalopathy and brain malformations [8–11]. Although utility is compromised by the comparative lack of reagents, the rat has many advantages over the mouse for the study of development, physiology, pathology and mononuclear phagocyte homeostasis (reviewed in [12]). We previously generated and characterised *Csf1rko* rats as an alternative model of human CSF1R deficiency [13]. This model was initially established and analysed on a mixed genetic background to avoid the pre-weaning mortality seen in *Csf1rko* mice. Although the gross osteopetrotic phenotype of these rats was 100% penetrant, post-weaning survival was variable and apparently female-biased [13]. To enable us to test complementation of the *Csf1rko* mutation by transfer of bone marrow (BM) and to establish a more consistent model we back-crossed the original outbred line to the dark agouti (DA) inbred background, which was the origin of the ES cells used in homologous recombination. Unlike inbred *Csf1rko* mice, the majority of the inbred DA *Csf1rko* rats survive to adulthood. In the present study we use the inbred line to analyse the profound impacts of the *Csf1rko* on tissue monocyte and macrophage populations and the growth and development of the skeleton and major organs in the early postnatal period. We show that CSF1R-dependent macrophages are essential in the liver for postnatal hyperplasia, hepatocyte functional maturation and lipid metabolism and that resident macrophage populations and the pleiotropic *Csf1rko* phenotypes can be almost completely reversed by transfer of BM cells from WT congenic animals at weaning. Our findings demonstrate that expansion and maturation of resident tissue macrophage populations is a key event in postnatal development that controls somatic growth and organ development.

## Results

### The effect of *Csf1rko* on embryo survival and postnatal growth in rats

Previous analysis of the *Csf1rko* on the outbred background focussed on surviving adult female animals [13]. The postnatal mortality was variable and apparently more penetrant in males. The impact of the mutation on the inbred DA background remained distinct from the pre-weaning lethality reported in inbred *Csf1rko* mice [3, 14, 15]. There was some evidence of early postnatal mortality, especially in first litters, but the large majority of *Csf1rko* pups survived. The genotype frequencies at weaning from a heterozygous mating were wild-type 0.27, Heterozygote 0.53, *Csf1rko* 0.19 based upon 65 litters. The majority of homozygous DA *Csf1rko* animals that survived to weaning were sacrificed because of breathing difficulties by 10 weeks.

To dissect the mechanisms underlying the *Csf1rko* phenotype, we first examined the effect of the *Csf1rko* on macrophage populations in the embryo. IBA1 staining labelled the abundant macrophage populations in the liver and throughout the WT embryos at around 10.5 days of gestation whereas staining was almost absent in *Csf1rko* embryos in the same litters (**Figure 1A, B)**. A similar lack of IBA1^+^ cells was observed at later gestational age (12.5-13.5dpc; not shown). One of the major functions of macrophages in the embryo is the clearance of apoptotic cells [16]. To examine this further we performed TUNEL (terminal transferase-mediated dUTP-biotin nick end-labelling) staining to detect apoptotic cells. There was no evidence of accumulation of TUNEL^+^ staining except for occasional cells in the liver (**Figure 1B, C**). Locations where CSF1R^+^ macrophages are actively involved in phagocytosis of apoptotic cells include the interdigital region and the pharyngeal arches [17]. However, we saw no evidence of delay in clearance of dying cells (e.g. accumulation of pyknotic nuclei) in these regions despite the complete absence of IBA1^+^ cells in the *Csf1rko* embryos (**Figure 1B**).

**Figure 1.**
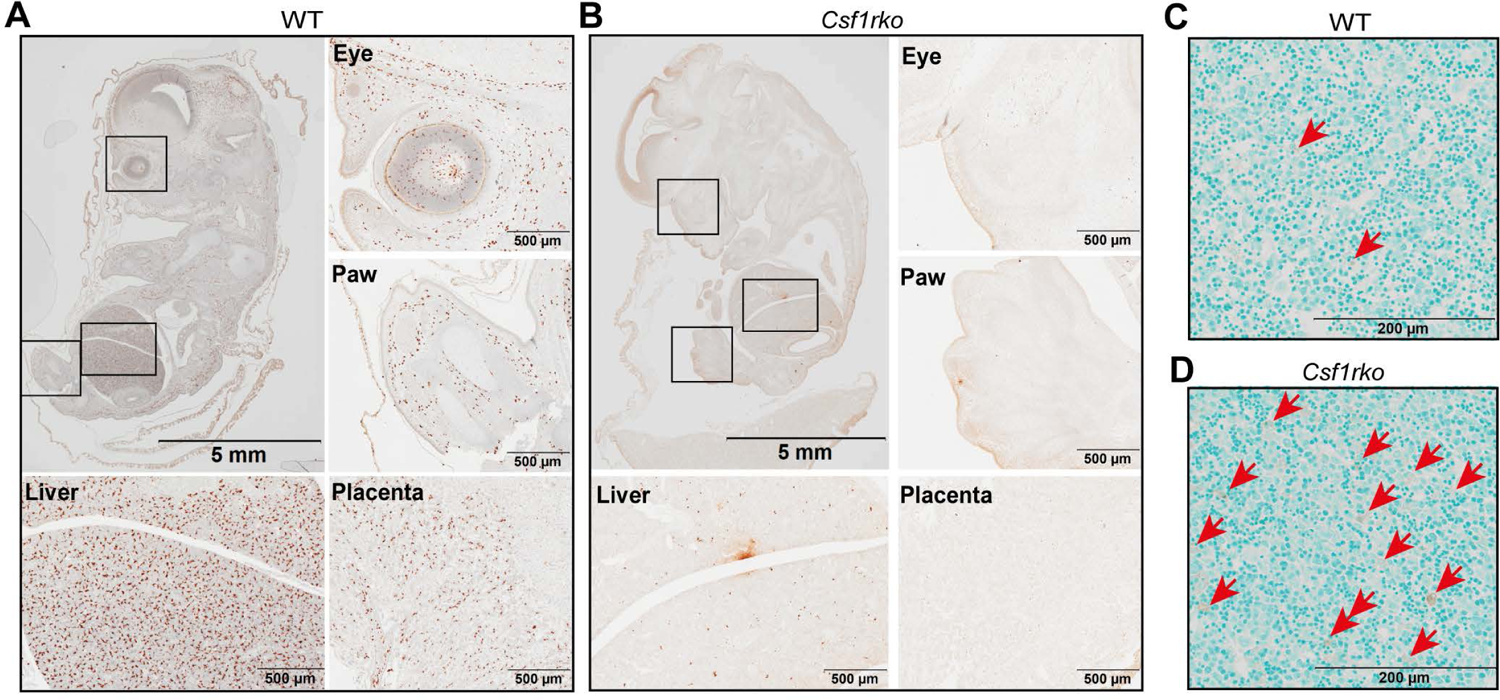
IBA1 expression and TUNEL staining in WT and *Csf1rko* embryos. (A, B) Histological analysis of IBA1 expression in embryos of (A) WT and (B) *Csf1rko* from the same litter approximately 10.5 days post conception (dpc). IBA1+ cells are detectable throughout the WT embryo and absent from the *Csf1rko.* The fetal liver, eye and paw (squares) and placenta is shown at higher magnification in the lower panels. Note that early digit condensation is evident in the hind limb of the *Csf1rko* embryo but there are no infiltrating IBA1^+^ cells. (C, D) TUNEL staining in the liver of (C) WT and (D) *Csf1rko* embryos approximately 10.5 dpc. TUNEL^+^ expelled nuclei retained in the *Csf1rko* liver are highlighted by red arrows.

Like the outbred animals, the inbred *Csf1rko* rats were indistinguishable from littermates at birth in either body weight or morphology. By 3 wks they were less than 50% of the weight of wild-type (WT) controls. Most organs, including the liver, were proportionally reduced but the absolute size of the brain was unaffected **(Figure S1A, B**). The relative lack of effect of the *Csf1rko* on the adult brain in outbred rats, aside from lateral ventricular enlargement, was described previously [13]. The very limited impacts of the *Csf1rko* on postnatal development of multiple brain regions on the inbred background are reported elsewhere [18].

As reported previously, the relative growth advantage of males over females was also abolished **(Figure S1C)**. By 7 wks of age, the shape of the brain was clearly different between WT and *Csf1rko* rats consistent with the radical difference in skull shape **(Figure S1D-F**). The reduced somatic growth was associated with significant reductions in cellular proliferation in major organs such as liver, lung and kidney as detected by localisation of Ki67 (**Figure 2A-F)**.

**Figure 2.**
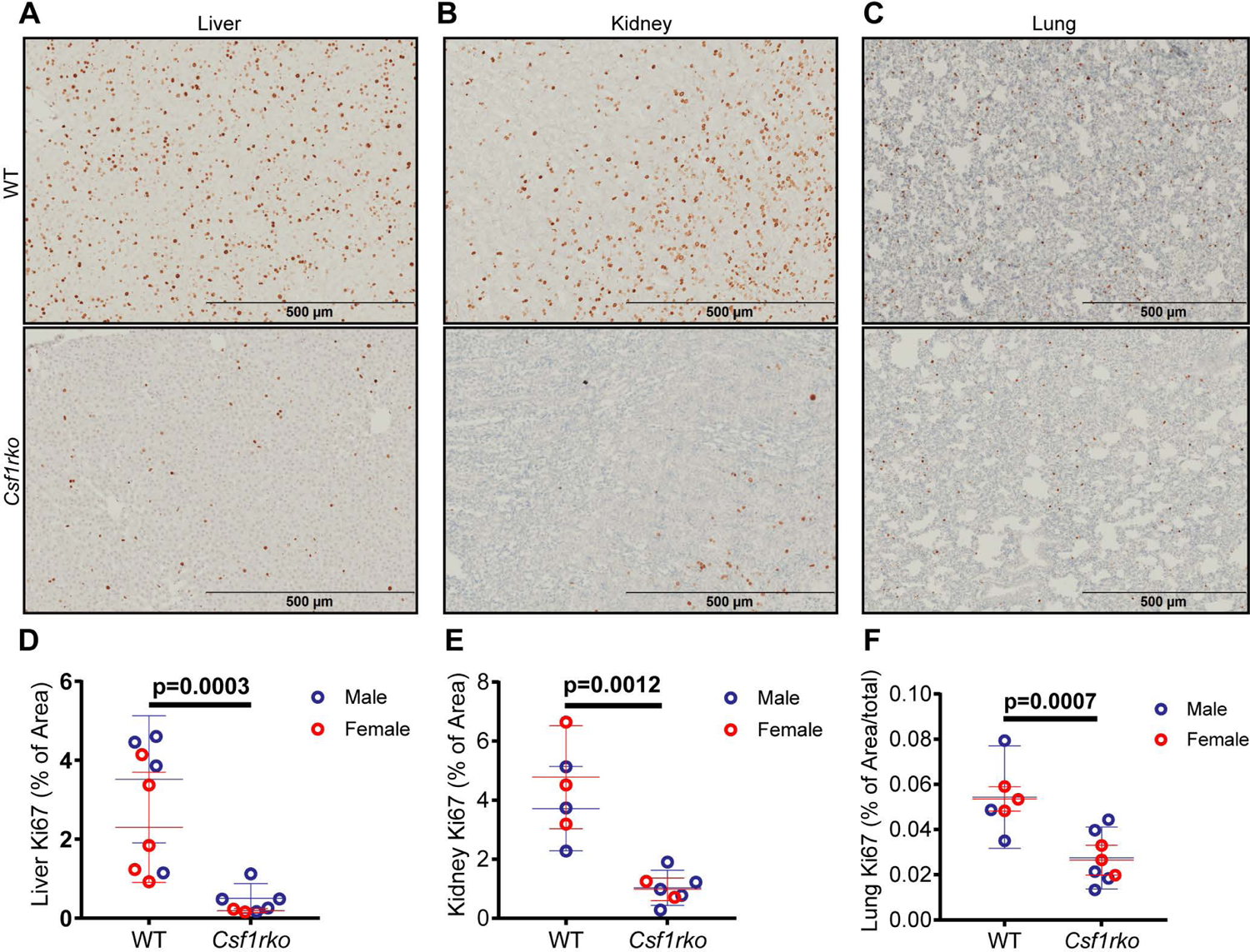
Detection of Ki67^+^ proliferating cells in the liver, kidney and lung of WT and *Csf1rko* rats at 3 wks of age. (A-C) Representative images of Ki67 staining in (A) liver, (B) kidney and (C) lung of WT and *Csf1rko* rats. D-F) Quantitative analysis of Ki67+ cells in (D) liver, (E) kidney and (F) lung. n≥6 per genotype, graphs shows the mean ± SD, genotype comparisons were analysed by Mann Whitney test.

### The effect of the *Csf1rko* on skeletal development

The skeletal phenotype of the outbred adult *Csf1rko* rat [13] closely resembles that of human CSF1R-deficient patients, referred to as dysosteosclerosis [8]. As shown in **Figure 3A**, analysis of the juvenile inbred *Csf1rko* rats revealed that a delay in skeletal calcification was already evident in newborns. By 3 wks of age secondary ossification centers of long bones and the small bones in the fore and hind paws were clearly deficient (**Figure 3B, C**). Unlike the hyper-calcified skull base [13], a shared feature with human bi-allelic *CSF1R* mutation [8, 10, 19], the cranial case of the *Csf1rko* rats remained hypomineralized even in adults and closure of the sutures was impaired (**Figure 3D**). The delay in postnatal musculoskeletal development was reflected in muscle. Sections through muscle at 7 days, 3 wks and 7 wks stained with laminin (**Figure 3E, F)** demonstrate that the relative cellularity was similar and the reduced muscle mass in the *Csf1rko* was primarily associated with a reduction in muscle fibre diameter (i.e. failure of hypertrophy).

**Figure 3.**
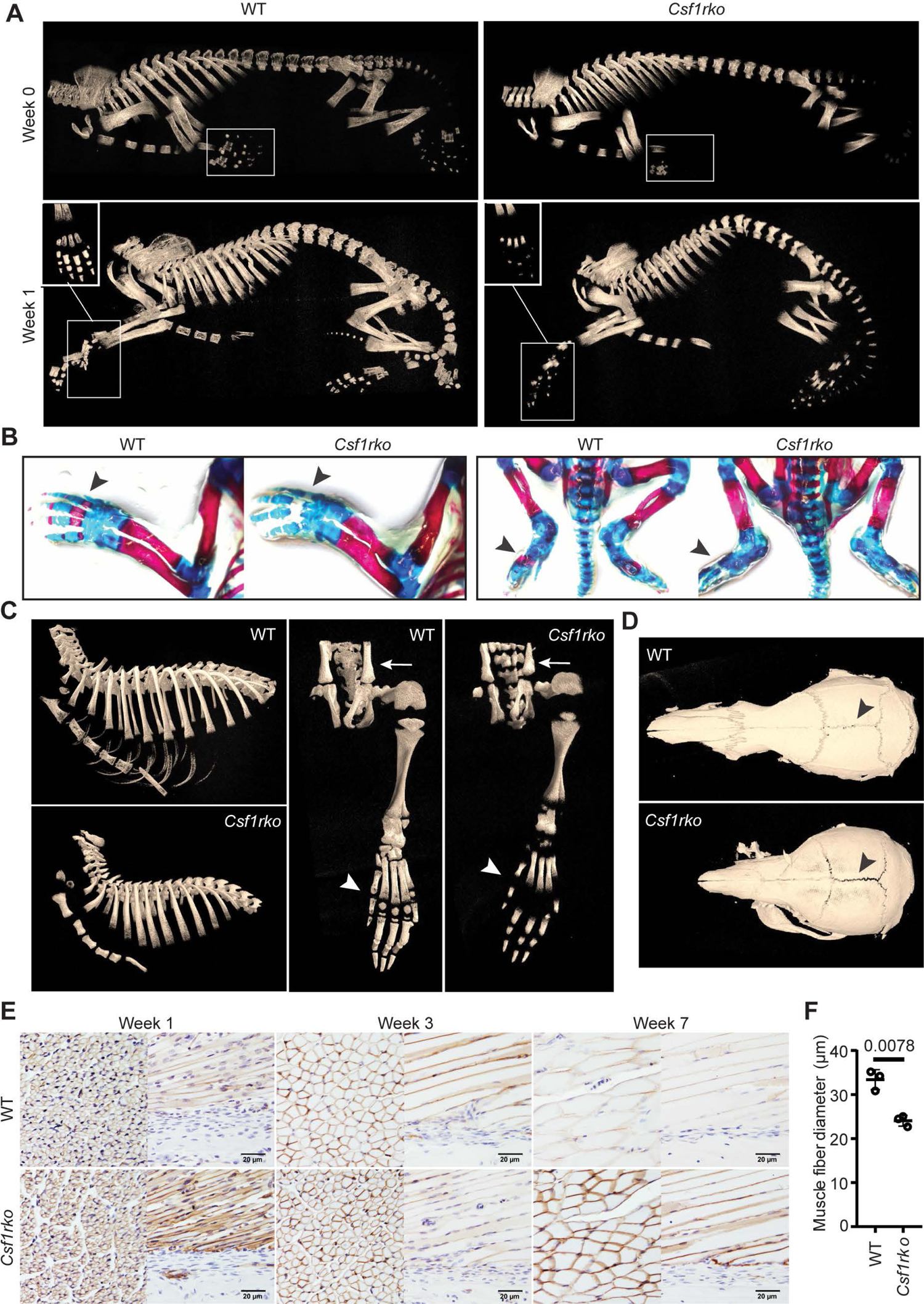
Postnatal skeletal development in *Csf1rko* rats. (A) Representative 3D reconstruction of micro-CT images in newborn (Week 0) and 1wk old WT and *Csf1rko* rats. Digits of forepaws in white boxes. (B) Alcian blue and Alizarin red staining of newborn WT and *Csf1rko* rats with ossified cartilage/bone stained red (black arrows) and unossified cartilage/bone stained blue. (C) Representative 3D reconstructed micro-CT images of 3-week old rat ribcage (left panel) and hind paw (right panel) with arrow heads indicating variance in ossification in distal metatarsal/proximal phalanges. (D) Representative 3D reconstructed micro-CT images of 7-wk old rat skull of WT (top) and *Csf1rko* (bottom) rats. (E) Representative images of laminin staining of hindlimb muscles of WT and *Csf1rko* rats at 1, 3 and 7 wks. (F) Average diameter of muscle fibres from the posterior tibialis in WT and *Csf1rko* rats at 3 wks. Original magnification: 400X. Scale bar: 20 µm.

The impact of the *Csf1rko* on the liver.

There is a well-known homeostatic relationship between liver and body weight [20, 21]. In the outbred adult *Csf1rko* rats analysed previously there was approximately 70% reduction in Kupffer cells (KC) detected in the liver, associated with selective loss of KC-enriched transcripts detected using microarrays [22]. To our knowledge there has been no previous analysis of the role of macrophages in postnatal liver development. In rodents, relative liver mass increases several fold in the postnatal period reaching the adult liver/body weight (LBW) ratio by around 4 wks of age [23, 24]. The postnatal proliferative expansion of the liver is associated with profound changes in gene expression. The FANTOM5 consortium generated a dense developmental time course of RNA expression in mouse liver using cap analysis of gene expression (CAGE). Network analysis of these data revealed clusters of co-expressed genes with distinct temporal profiles including postnatal expansion of a macrophage-related cluster [25]. To assess the role of CSF1R in this hepatic differentiation/maturation process in the rat, we generated expression profiles of the inbred male and female *Csf1rko* and WT rats at 3 wks of age by RNA-seq. **Table S1A** contains the primary data and **Table S1B** shows 2760 differentially-expressed genes (DEG) distinguishing WT and *Csf1rko* livers (FDR <0.05). The DEG were grouped into a number of categories based upon known functions (i) growth factors (ii) Kupffer cell/macrophage-associated (iii) cell cycle-related; (iv) lipid metabolism (v) liver function/maturation-associated. **Figure 4A** summarises representative DEG in each category. The genes significantly reduced in the *Csf1rko* included *Ghr, Igf1* and *Igfals,* with the latter most affected. *Igf1* and *Ghr* closely paralleled each other in individual WT and *Csf1rko* samples **(Table S1A**). By contrast, the genes encoding the major IGF1 inhibitory binding proteins *Igfbp1* and *Igfbp2* (which in mice were massively induced at birth and declined rapidly thereafter; [25]) were highly-expressed and amongst the most over-expressed transcripts in the 3 wk-old *Csf1rko* liver relative to WT. Aside from *Igf1*, known direct GH targets (e.g. *Socs2, Socs3, Cish;* [26]) were unaffected suggesting that GH signalling was not impaired.

**Figure 4.**
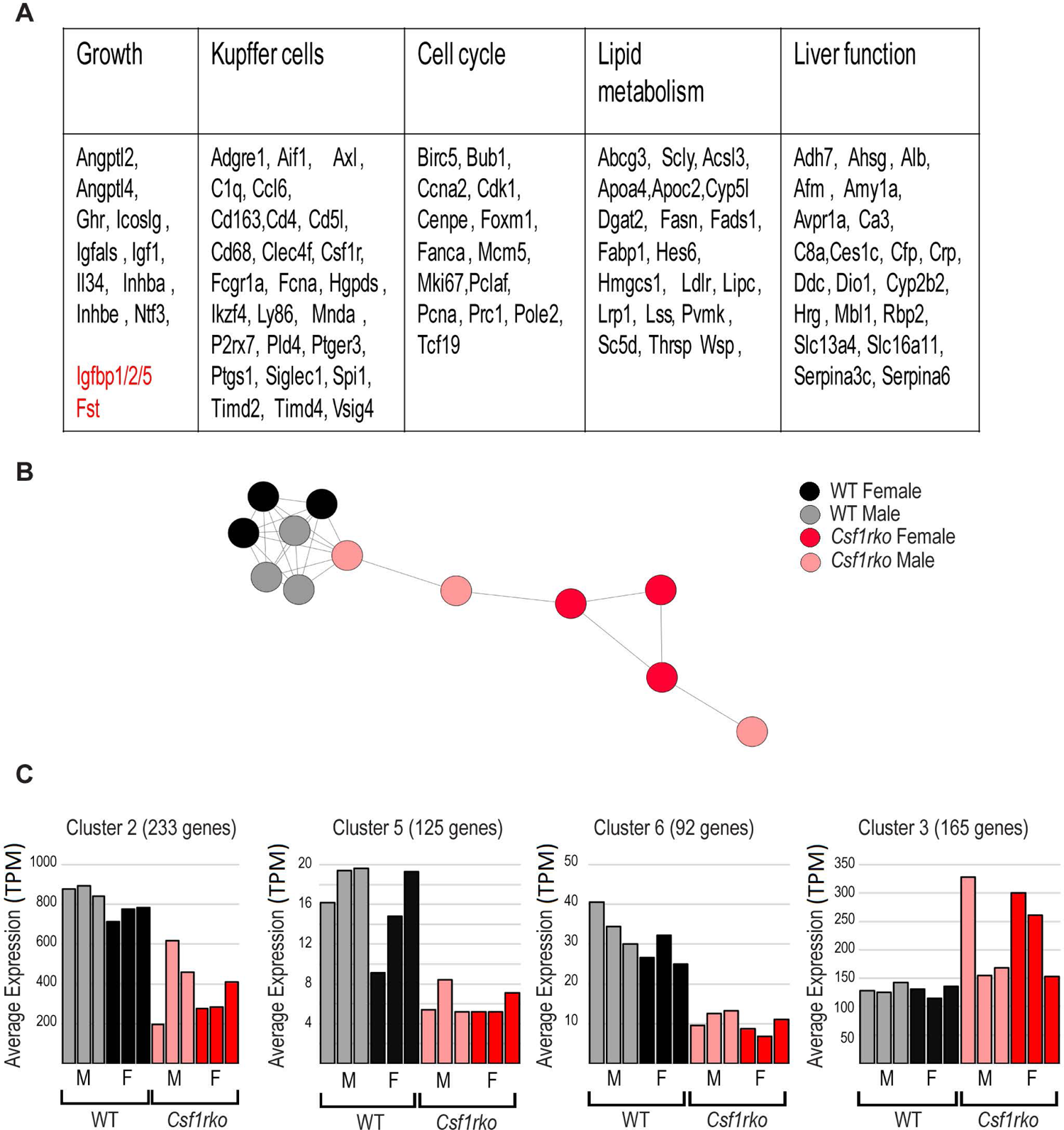
Analysis of gene expression in livers of WT and *Csf1rko* rats. RNA-seq analysis was performed on liver from 3 male and 3 female wild-type and *Csf1rko* rats at 3 wks of age. The full data set and the differentially-expressed gene analysis are provided in Table S1A and S1B. Co-expression cluster analysis was performed using BioLayout as described in Materials and Methods. (A) Summary of regulated genes. Genes shown were regulated >2 fold in the livers of *Csf1rko* rats compared to WT. They have been assigned to categories based upon known function or expression. Genes in red encode proteins that inhibit liver or somatic growth and are over-expressed in the *Csf1rko*. (B) Sample to sample analysis of *Csf1rko* and wild type live samples. Circles represent samples and lines between them show correlations of r≥0.93 between expression patterns of pairs of samples across all genes. (C) Average expression profile of four clusters of co-expressed transcripts generated using BioLayout that distinguish *Csf1rko* and wild-type liver. Full list of transcripts within each cluster is provided in Table S1C and GO annotation in Table S1D. Y axis shows the average expression of transcripts in the cluster (TPM); X axis shows the samples with columns colored as for (B).

To identify gene co-expression networks, we analysed the data using the analysis and visualisation tool BioLayout (http://biolayout.org). The sample-to-sample network graph for all transcripts revealed a clear separation based upon genotype but a lack of separation based upon sex (**Figure 4B)**. Clusters of co-expressed transcripts identified from a gene-to-gene correlation network are summarised in **Table S1C** and gene ontology (GO) terms for the largest clusters in **Table S1D. Figure 4C** shows the average profiles of the 4 co-regulated gene clusters that distinguished the WT and *Csf1rko*. The three clusters with lower average expression in the *Csf1rko* are consistent with the relative loss of Kupffer cells (Cluster 6), reduced proliferation and cell cycle (Cluster 5) and delayed liver-specific differentiation (Cluster 2). Cluster 3, which contains transcripts expressed more highly, but variably, in the *Csf1rko* compared to WT is enriched for transcripts associated with mitochondria and oxidative phosphorylation.

As observed previously in outbred juvenile WT rats [27] the livers of both WT and *Csf1rko* DA rats were mildly steatotic at 3 wks of age. Whereas this resolved with age in WT there was progressive and extensive steatosis in the liver of older *Csf1rko* rats (**Figure 5A)** which was not noted previously. Because of the failure of tooth eruption, the *Csf1rko* rats were maintained on a modified diet (see Methods) but steatosis was not observed in the livers of WT rats maintained for 12 wks with access to the same diet (not shown). The *Csf1*^op/op^ mouse has been reported to have a substantial deficit in insulin-producing cells in the pancreatic islets [28]. The selective loss of visceral adipose tissue [27], which was also evident in the inbred line, might suggest the reverse in the *Csf1rko* rat. We therefore measured circulating fed glucose and insulin levels and found small but significant reduction in fed insulin and glucose levels in the *Csf1rko* at 7 wks of age (**Figure 5B, C**).

**Figure 5.**
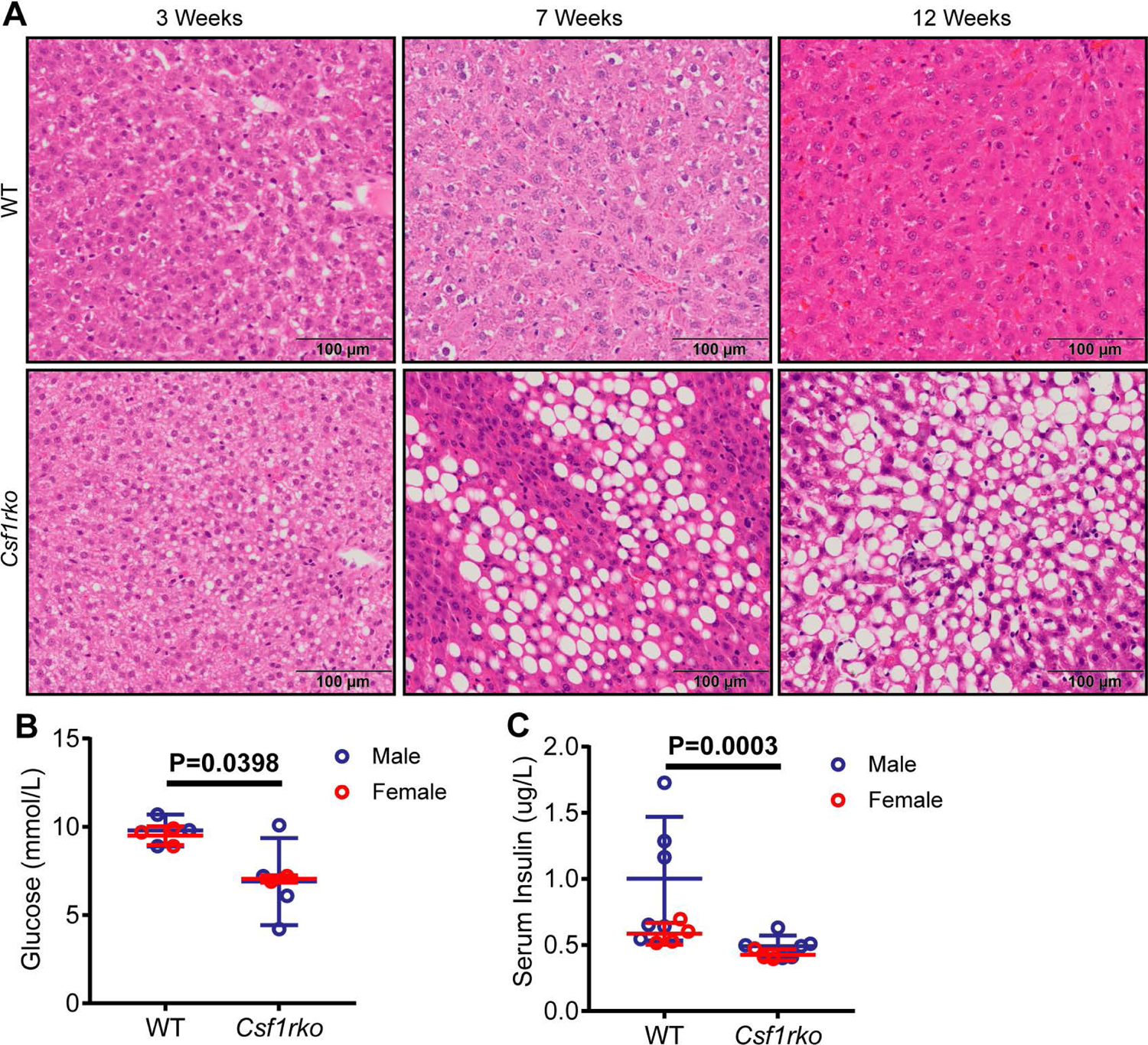
Histological analysis of lipid accumulation in the liver and quantitative measures of blood glucose and serum insulin. (A) Representative images of H&E staining in the liver of WT (top panel) and *Csf1rko* (bottom panel) rats at 3 wks (left panels), 7 wks (middle panels) and 12 wks (right panels). (B) Blood glucose and (C) serum insulin levels measured at 3 wks in WT and *Csf1rko* rats. n≥6 per genotype, graphs show the mean ± SD, genotype comparisons were analysed by Mann Whitney test.

### The effect of the *Csf1rko* on major organs

Further analysis of juvenile rats indicated that the developmental delay in the inbred *Csf1rko* was not confined to the liver. One phenotype we did not note previously was almost complete involution of the thymus which was almost undetectable by 7 weeks of age (not shown). A subset of major organs is shown in **Figure S2**. In the rat, the process of nephrogenesis continues in the immediate postnatal period [29]; the impact of the *Csf1rko* on the kidney was not previously analysed [13]. By 7 wks of age, we observed profound renal medullary hypoplasia, which was so severe that in most *Csf1rko* rats examined there was just a bud of papilla within the central pelvicalyceal space. The cortex was correspondingly hyperplastic, but the tubules and glomeruli appeared relatively normal **(Figure S2A,B**).

The intestines of outbred adult *Csf1rko* rats were not grossly abnormal and by contrast to mice with the ligand mutation (*Csf1*^op/op^), they were not Paneth cell deficient [13]. To dissect more subtle impacts of the *Csf1rko* in the inbred line we performed quantitative analysis of villus architecture in the ileum. There were significant reductions in the length and width of the crypts and villi, and the thickness of the submucosa in the *Csf1rko* **(Figure S2C-F)**. Interestingly, given the reported role of CSF1R dependent macrophages in M cell differentiation in mice [30], we noted for the first time that macroscopic Peyer’s patches were almost undetectable in the *Csf1rko* rats at 3 wks and remained so at later ages.

The main welfare concern with the inbred *Csf1rko* is the development of progressive breathing difficulty. We considered the possibility that macrophage deficiency might also impact postnatal development of the lung. **Figure S2G** shows images of the inflated lungs of WT and *Csf1rko* rats at 3 wks stained with aldehyde-fuchsin to highlight elastin fibres and **Figure S2H** quantitates the airway space. There was no unequivocal evidence of impaired alveolisation. Flow cytometric analysis of disaggregated lung tissue revealed an increase in granulocytes in *Csf1rko* rats by 9 wks (**Figure S2I, J**), as seen in peripheral blood and BM, but there remained no histological evidence of inflammation that could explain the impaired respiratory function.

As in the outbred line [13], male and female inbred *Csf1rko* rats lacked development of primary and secondary reproductive organs. In males, the prostate and seminal vesicles and in females the uterus were so under-developed as to be almost undetectable (not shown). CSF1R*-*dependent macrophages have also been implicated in the process of branching morphogenesis in the mammary gland in mice [31]. This was not previously analysed in the rat. **Figure S 3** shows a comparison of mammary gland development in *Csf1rko* and WT female DA rats at around 9 wks. The negative impact of the mutation on ductal development was evident from staining with mature epithelial cell markers (Krt5, E-cadherin).

### The effect of the *Csf1rko* on tissue macrophages

Aside from limited analysis using CD68 as a marker in adult liver and spleen, the effect of the *Csf1rko* on resident macrophages aside from microglia was not examined previously [13]. To visualise resident macrophages we crossed the *Csf1rko* back to the *Csf1r*-mApple reporter transgene on the outbred SD background [32]. **Figure 6** shows detection of the mApple transgene in a diverse set of tissues from WT and *Csf1rko* rats on this genetic background at weaning. In most tissues there was complete loss of *Csf1r*-mApple expressing cells aside from occasional monocyte-like cells and granulocytes in the vessels. This includes the abundant resident macrophage populations in smooth and skeletal muscle, kidney, pancreas, adipose, salivary and adrenal glands that were not recognised or analysed previously in mice. The pancreas contains numerous small lymph nodes that contain abundant *Csf1r-*mApple-positive cells. Whereas peri-acinar and islet macrophage populations were entirely lost in the *Csf1rko* the lymph node-associated populations were partly retained and highlighted in whole mounts (**Figure 6D**). Partial reductions in resident populations were observed in intestinal mucosa, liver and lung (**Figure 6C, 6H, 6J**) Note that the resident *Csf1r*-mApple expressing cells in non-lymphoid tissues that are missing in the *Csf11rko* would include populations classified as dendritic cells, which are also CSF1R-dependent in mice [33].

**Figure 6.**
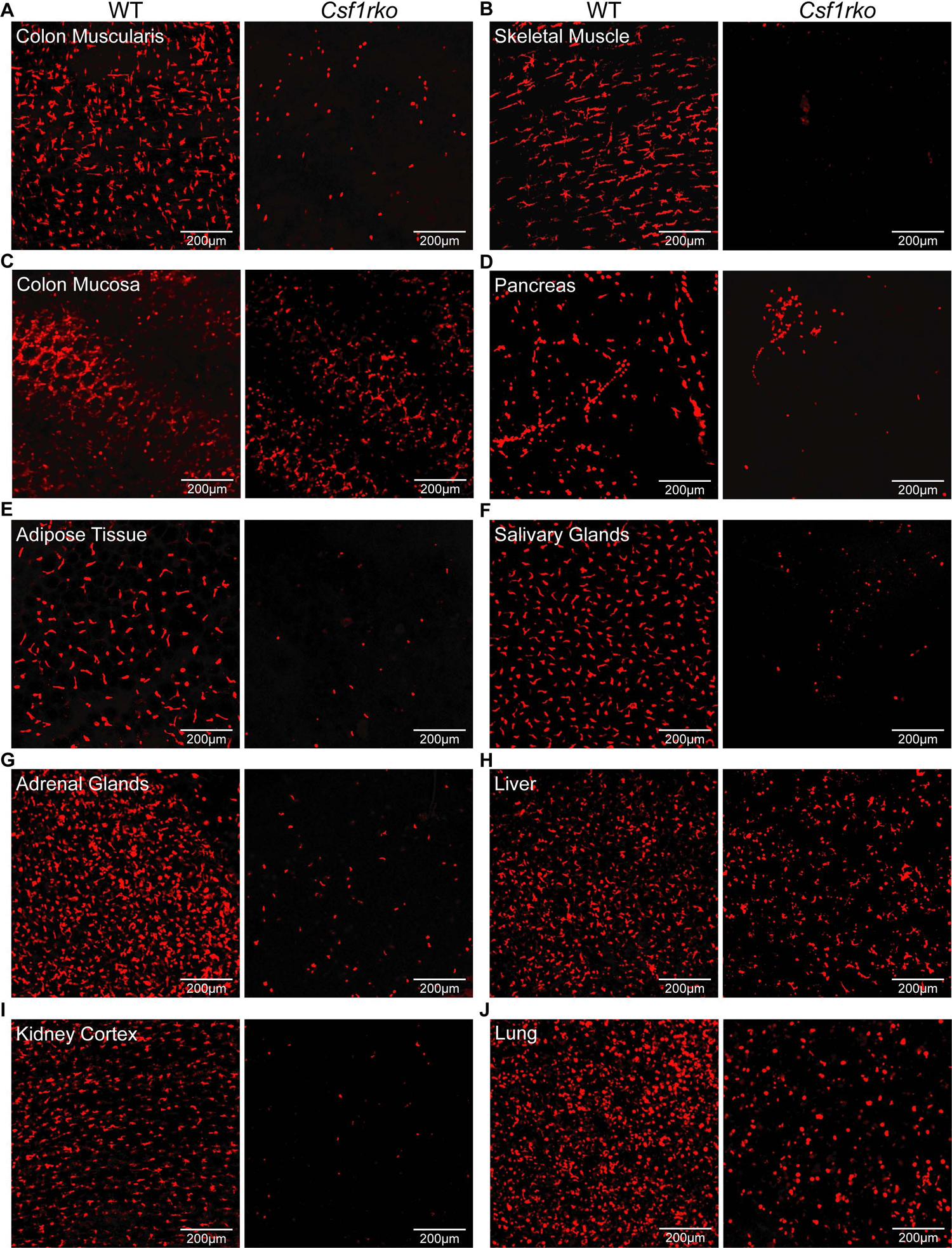
The effect of the *Csf1rko* on distribution of macrophages detected using a *Csf1r-*mApple reporter gene. Tissues were extracted from 6-7 week old rats generated by crossing heterozygous inbred DA *Csf1rko* rats with outbred (SD) *Csf1r-*mApple reporter rats and then inter-breeding selected progeny to generate wild-type and mutant rats also expressing the mApple reporter. Images are maximum intensity projection of z-stack series of whole-mounted (A) colon muscularis (smooth muscle), (B) skeletal muscle, (C) colonic mucosa, (D) pancreas, (E) perigonadal white adipose tissue, (F) salivary glands, (G) adrenal glands, (H) liver, (I) kidney cortex and (J) lung. Scale bars = 200um.

Rescue of the *Csf1rko* phenotype by wild type bone marrow cells and the role of IGF1.

To determine whether the major developmental abnormalities in the *Csf1rko* rat were autonomous to hematopoietic lineage cells and reversible we treated a cohort of 3 wk-old DA *Csf1rko* rats with WT congenic BM cells by IP injection without ablation of recipient BM. By 2-3 wks post BM transfer the body weight gain of recipients diverged from *Csf1rko* controls and continued to increase thereafter (**Figure 7A**). The recipients did not fully recover the deficit in growth. At necropsy up to 6 months post BM transfer, the body weights were around 10% lower than littermates (**Figure 7A**) but visceral adipose was fully restored in both sexes and major organ histology was indistinguishable from controls.

**Figure 7.**
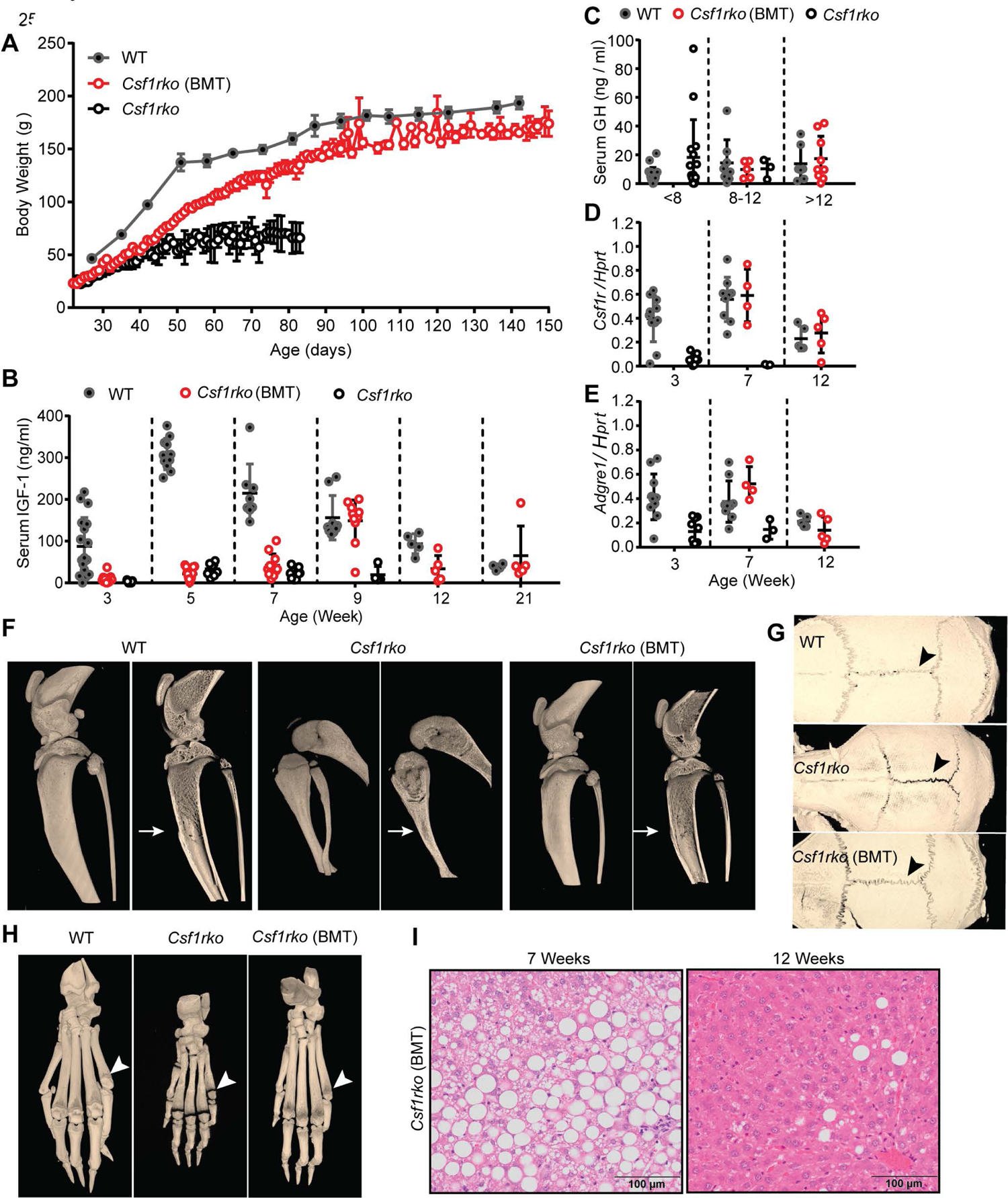
Rescue of the *Csf1rko* phenotype by transfer of WT bone marrow. A cohort of 9 female and 2 male inbred *Csf1rko* rats received WT bone marrow cells by intraperitoneal injection at 3 wks of age. (A) Time course of body weight gain in female WT (n≥3) *Csf1rko* rats with WT bone marrow transfer (BMT) and *Csf1rko* from 3 wks of age to harvest. Note that the *Csf1rko* control rats in this cohort were euthanised at 11-12 wks for welfare reasons. (B) Serum IGF-1 levels and (C) serum growth hormone (GH) levels in a mixed cohort of male and female WT, *Csf1rko* following bone marrow transfer (BMT) at 3 wks and *Csf1rko* rats at the ages indicated. (D, E) qRT-PCR analysis of liver (D) *Csf1r* and (E) *Adgre1* expression in 3-, 7- and 12-wks old WT, *Csf1rko* (BMT) *and Csf1rko* rats. n≥3 per genotype, graphs show the mean ± SD. (F-H) Representative 3D reconstructed micro-CT images of WT, *Csf1rko* and *Csf1rko* (BMT) rat (F) hind limbs, (G) hind paws and (H) skulls comparing medulla cavity (arrows), ossification of phalanges (white arrowheads) and closure of cranial sutures (black arrowheads). (I) representative images of H&E staining in the liver at 7 and 12 wks of age of *Csf1rko* (BMT) rats after WT bone marrow transfer.

The appearance of the animals changed rapidly, notably the skull and limb/foot morphology, so that by 6 wks post-transplant they resembled WT animals (**Figure 7F-H**). The steatosis observed in untreated *Csf1rko* rats remained 4 wks post BM transfer but resolved by 9 wks (**Figure 7I**). Many BMT recipients developed teeth which required regular trimming. Both males and females became sexually mature and fertile and produced multiple litters. The females were able to suckle their offspring indicating that the failure of mammary development was reversible as confirmed in **Figure S3**.

The *Csf1*^tl/tl^ rat has been reported to have a deficiency in circulating IGF1 [34] and the transcriptomic analysis of the liver indicated dysregulation of the GHR/IGF1/IGFBP system. Accordingly, we measured IGF1 and GH in the circulation of the *Csf1rko* rats. In the WT DA rats, there was a postweaning surge in IGF1 followed by a gradual decline (**Figure 7B)** whereas IGF1 was barely detectable in the serum of *Csf1rko* DA rats at any age. By contrast, the circulating concentration of GH was unaffected (**Figure 7C)**. The DA rat is relatively small and slow-growing compared to outbred lines such as Sprague-Dawley (SD). To test the impact of genetic background, we crossed the mutation back out to SD for two generations. Despite the more rapid postnatal growth of SD and 2-3 fold higher adult body weight, both the time course and magnitude of circulating IGF1 was very similar between the WT inbred and outbred lines. Unlike the almost complete loss of IGF1 seen in the DA *Csf1rko* line, the surge of circulating IGF1 in the postnatal period was readily detected, albeit reduced and delayed in the SD *Csf1rko* rats **(Figure S4A)**. Nevertheless, the SD *Csf1rko* rats showed the same growth arrest around 7 weeks of age as the DA **(Figure S4B)**, albeit at peak body weight of 80-100g rather than 50-70g. In the DA *Csf1rko* rats that received WT BM, IGF1 was only partly restored. It was first detectable by 4 wks post transfer (week 7) and peaked at 6 wks (week 9) (**Figure 7B**). However, the divergence of body weight gain in the BMT recipients compared to untreated *Csf1rko* rats was evident within 2 weeks, before IGF1 was detectable and peak levels did not recapitulate the postnatal surge.

### Recovery of tissue macrophage populations

The *Csf1r-*mApple transgene is not currently available on an inbred background to enable analysis of tissue macrophage recovery following BMT. For this purpose, as in the embryo (**Figure 1**), we used IBA1 as a marker. Although most commonly used as a microglial marker, IBA1 is widely expressed by tissue macrophages in mice [35]. Immunohistochemical localisation of IBA1 in selected organs from the inbred *Csf1rko* and litter mate control rats at 3 and 7 wks is shown in **Figures 8 and 9**. In most WT tissues, IBA1 immunoreactivity was restricted to abundant interstitial stellate cells resembling macrophages. The morphology, abundance and location of IBA1^+^ populations was comparable to the *Csf1r*-mApple transgene (**Figure 6)** and much more extensive than seen with anti-CD68 (ED1) [13]. The *Csf1rko* led to substantial or complete loss of macrophage-like IBA1**^+^** cells in all organs examined and there was no evidence of recovery with age.

**Figure 8.**
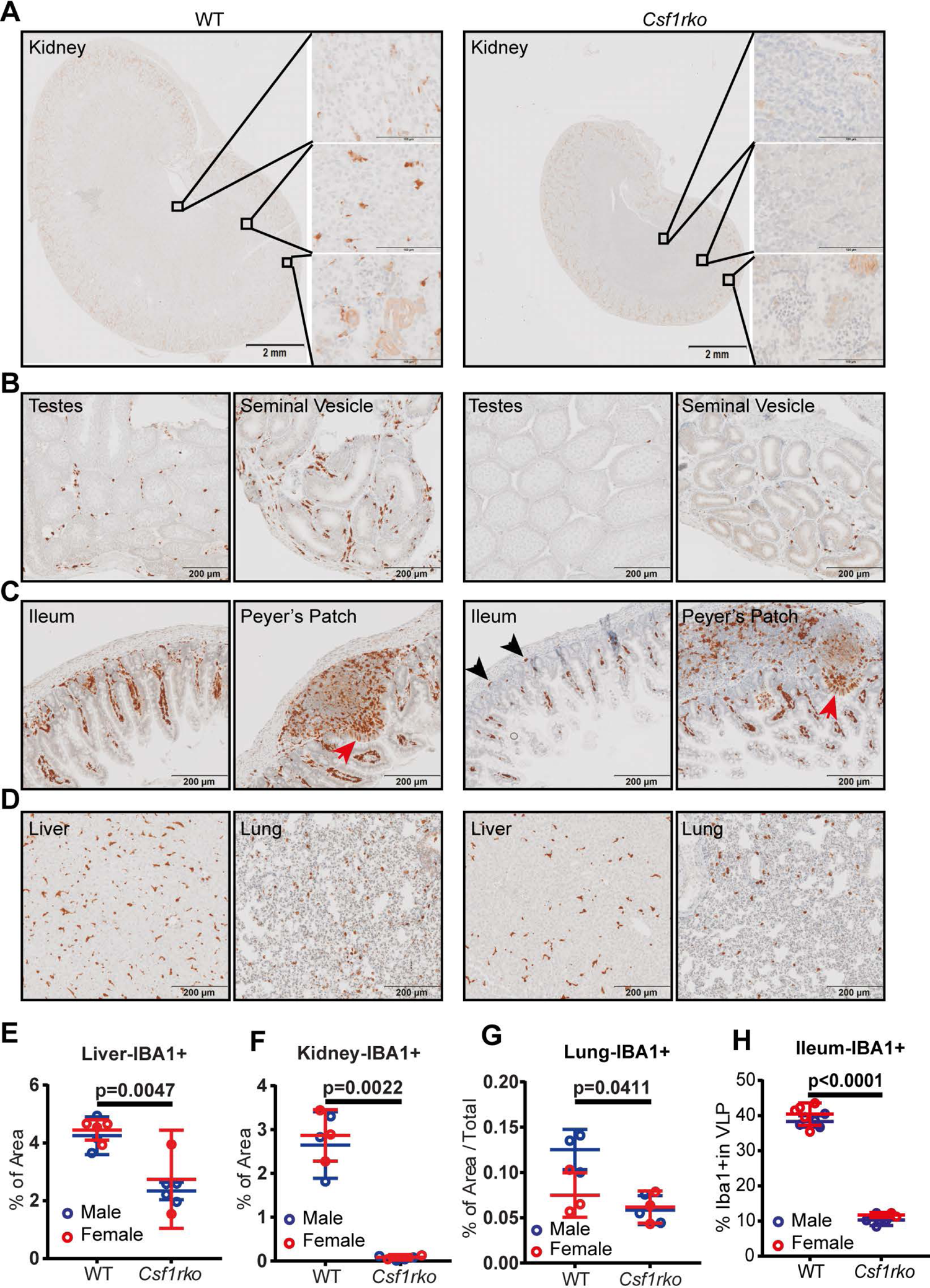
Immunolocalization of IBA1^+^ cells in WT and *Csf1rko* rat organs at 3 wks. IBA1 was detected in rat tissues by immunohistochemistry as described in Methods. Brown stain (diaminobenzidine) is a positive signal; sections were counterstained with haematoxylin (Blue). (A-D) Representative images of immunohistochemical localisation of IBA1 in (A) kidney, (B) testes and seminal vesicles, (C) ileum including Peyer’s patches and (D) liver and lung of WT and *Csf1rko* rats. M cells in the follicle-associated epithelium are indicated by red arrows. Black arrowheads highlight IBA1^+^ cells in the base of the crypts in (C). (E-G) Quantitative analysis of IBA1+ cells in (E) liver, (F) kidney and (G) lung. n ≥ 6 per genotype, graphs show the mean ± SD, ≥ genotype comparisons were analysed by Mann Whitney test.

**Figure 9.**
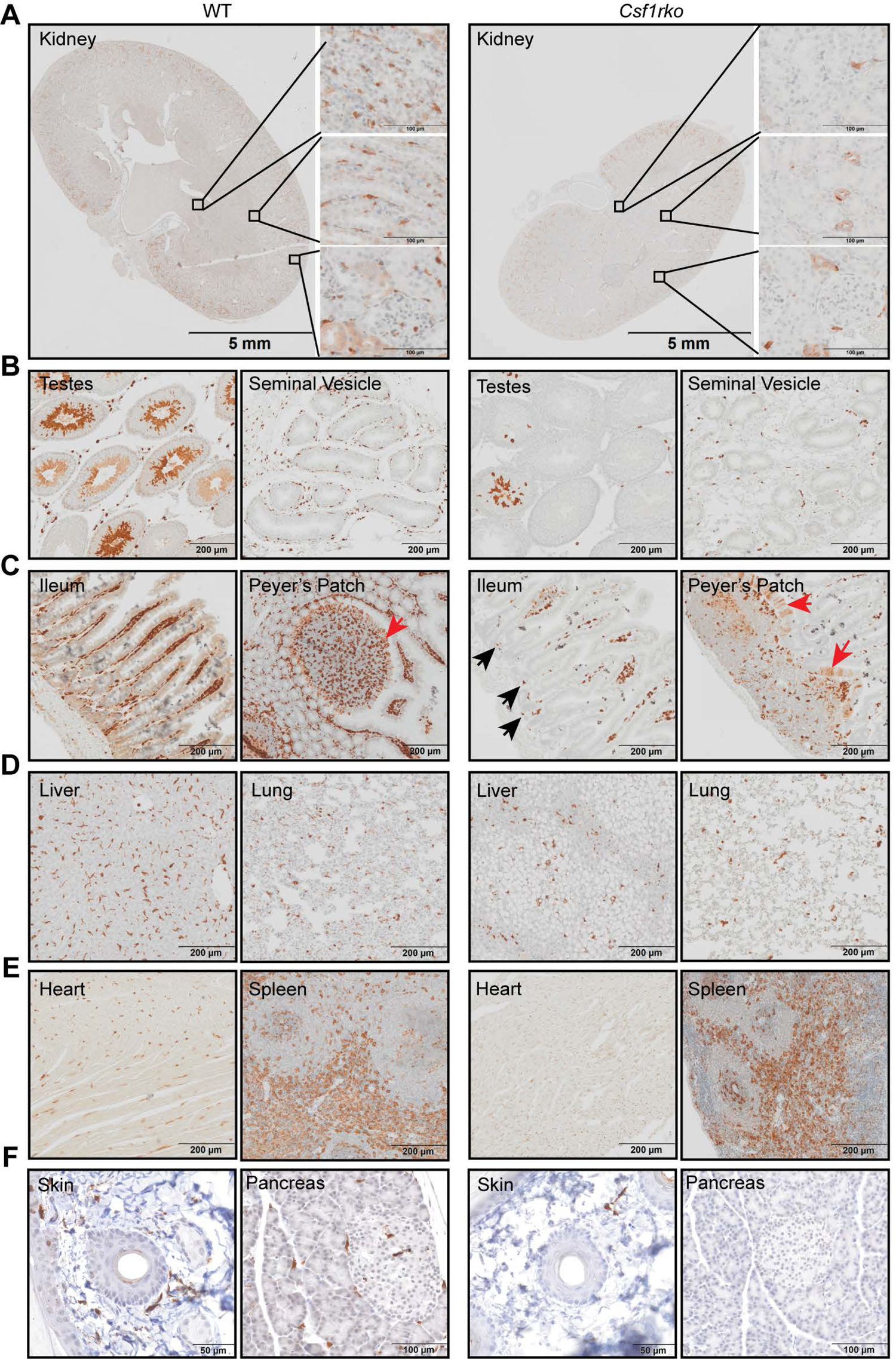
Immunolocalization of IBA1^+^ cells in WT and *Csf1rko* rat organs at 7 wks. IBA1 was detected in rat tissues by immunohistochemistry as described in Methods. Brown stain (diaminobenzidine) is a positive signal; sections were counterstained with haematoxylin (Blue). Representative images show localisation of IBA1 in (A) kidney, (B) testes and seminal vesicles, (C) small intestine including Peyer’s patches, (D) lung and heart and (E) liver and spleen of WT and *Csf1rko* rats. (F) Tail skin and (G) pancreas. M cells in small intestine are indicated by red arrows in (C). Black arrowheads highlight IBA1^+^ cells in the base of the crypts in (C).

The majority of IBA1^+^ tissue macrophage populations were restored by BMT. Resident peritoneal macrophages which were absent in the *Csf1rko* were restored to the level of WT controls. **Figure 10 A** shows IBA1 staining of multiple organs of the recipients of WT bone marrow indicating the restoration of tissue IBA1^+^ cell populations to levels similar to littermate controls. Note in particular that the visceral adipose which was restored following BMT, was populated with IBA1^+^ macrophages. BMT prevented premature thymic involution and the thymus of BMT recipients contained abundant IBA1^+^ cells. IBA1 is also expressed during sperm maturation, and IBA staining highlights the loss of mature sperm in testis (**Figure 9**) and their restoration following BMT (**Figure 10**). The repopulation of the liver with macrophages of donor origin was confirmed by analysis of the restoration of the expression of *Csf1r* (**Figure 7D)** and *Adgre1* mRNA (**Figure 7E**).

**Figure 10.**
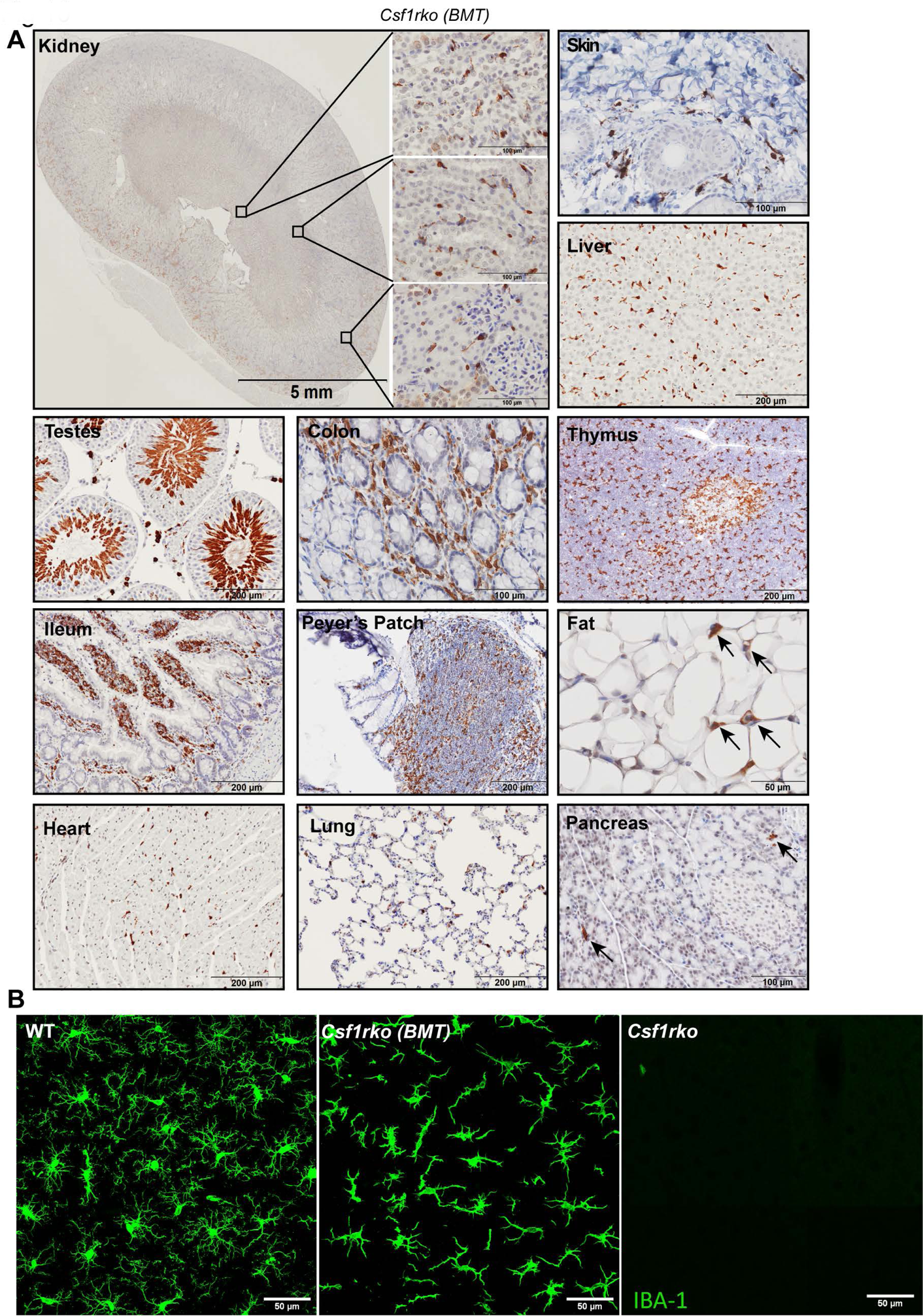
Immunolocalization of IBA1+ cells in *Csf1rko* rat organs after BM transfer IBA1 was detected in rat tissues by immunohistochemistry as described in Methods. Brown stain (diaminobenzidine) is a positive signal; sections were counterstained with haematoxylin (Blue). Inbred *Csf1rko* rats received WT bone marrow cells by intraperitoneal injection at 3 wks of age. (A) Representative images show immunohistochemical localisation of IBA1 in kidney, skin, liver, testes, colon, thymus, small intestine, Peyer’s patches, perigonadal fat pad, heart, lung, and pancreas. Note that BMT restores perigonadal visceral fat pads which are populated with interstitial IBA1+ cells (arrows). Image of pancreas contains exocrine (acinar) and endocrine (islet) regions. Note that only the exocrine region was partially repopulated with IBA1^+^ cells (arrows). (B) Confocal images of cortical microglia from 22-week-old male WT rat (left), *Csf1rko* rat of the same age which received BMT at 3 wks (middle) and 7-week-old *Csf1rko* rat (right), scale bar = 50 µm. The distinct stellate morphology in the BMT recipient is representative of all brain regions examined in 3 rats analysed.

The brains of *Csf1rko* rats lack microglia and brain-associated macrophages detected with anti-IBA1, alongside the loss of *Aif1* mRNA encoding this marker and many other microglia-associated transcripts [13, 18]. **Figure 10B** compares IBA1 staining in the cortex of adult WT, *Csf1rko* and BMT rats at 16 weeks post-BMT. The images are representative of all brain regions in all recipients examined. The IBA1^+^ cells in the BMT recipients adopted the same regular spacing and similar density to IBA1^+^ microglia detected in age-matched litter-mates. However, the cell morphology of the BMT-derived cells throughout the brain was quite distinct, more stellate and less ramified than typical microglia. The ventricular enlargement observed in juvenile *Csf1rko* rats was not reversed by BMT, but also had not progressed. The mechanism underlying hydrocephalus in both mouse and rat *Csf1rko* and bi-allelic human *CSF1R* mutations are unknown [6] but likely unrelated to microglial deficiency [36]. Otherwise, we could detect no difference between BMT recipient and WT littermate brains.

The concentration of CSF1 in the circulation is relatively low due to receptor-mediated clearance mainly by the macrophages of the liver and spleen [6]. Accordingly, increased circulating CSF1 was detected in the serum of adult *Csf1rko* rats by Western blot [13]. We reasoned that injected BM cells likely responded to the elevated CSF1 and successful restoration of CSF1R-expressing tissue macrophage populations could be monitored by measuring circulating CSF1 by ELISA. Indeed, CSF1 was massively elevated in juvenile *Csf1rko* rats at weaning and declined rapidly following BMT **(Figure S5A**). Given the granulocyte accumulation seen in the *Csf1rko* we also assayed CSF3 (granulocyte colony-stimulating factor) in the same samples, and there was no significant impact of the *Csf1rko* **(Figure S5B**).

In overview, *Csf1rko* rats are deficient in resident macrophages in most organs throughout postnatal development and rescue by BMT is associated with restoration of these populations to wild-type density.

The effect of the *Csf1rko* on blood and bone marrow mononuclear phagocyte populations.

*Csf1rko* mice were reported to have enlarged spleens and evidence of extramedullary hematopoiesis [37]. This is not the case in *Csf1rko* rats. Given the apparent lack of macrophages within hematopoietic islands in the fetal liver (**Figure 1**) we were especially interested in resident BM macrophages, which are believed to be an essential component of the hematopoietic niche [38]. As previously reported in outbred animals [13], the inbred *Csf1rko* rats were entirely deficient in osteoclasts expressing tartrate-resistant acid phosphatase (TRAP) (**Figure 11 A**). However, IBA1^+^ positive island macrophages were detectable in the residual BM cavity of *Csf1rko* rats at 7 weeks with similar stellate morphology to WT (**Figure 11A**).

**Figure 11.**
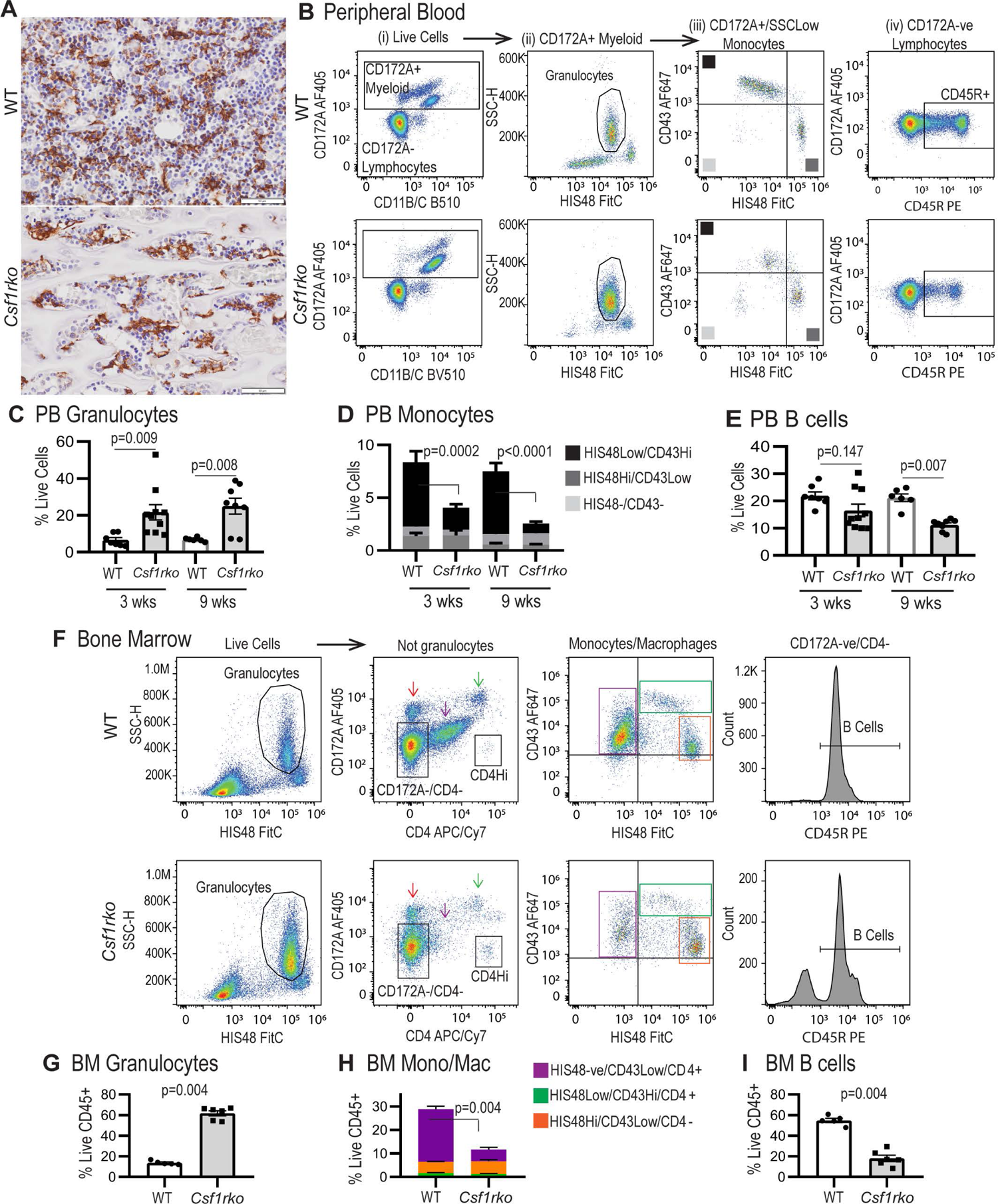
Flow cytometric analysis of peripheral blood and bone marrow leukocytes. (A) Representative images showing staining for IBA1 expression (brown) in naive 7-week-old WT and *Csf1rko* rat tibiae. Original magnification: 40X. Scale bar: 50µm. (B) Flow cytometry gating strategy to distinguish (i) CD172A+ myeloid cells, (ii) HIS48^+^/SSC^Hi^ granulocytes among myeloid cells, (iii) HIS48 and CD43-expressing monocyte subsets among non-granulocyte myeloid cells and (iv) CD45R^+^ B cells among CD172A^-^ lymphocytes. (C-E) Quantification of peripheral blood (C) granulocytes, (D) monocyte subsets and (E) B cells in WT and *Csf1rko* rats at 3 and 9 wks of age. (F) Bone marrow flow cytometry gating strategy to distinguish (i) HIS48^+^/SSC^Hi^ granulocytes, (ii) CD172^+^ and CD172^Low^/CD4^Int^ monocytes/macrophages among non-granulocytes, (iii) monocyte/macrophage subsets and (iv) CD45R+ B cells among CD172A-/CD4-cells. (G-I) Quantification of BM (G) granulocytes, (H) monocyte/macrophage subsets and (I) B cells. All genotype comparisons were analysed by Mann Whitney test.

Like adult outbred *Csf1rko* rats [13], juvenile (3 wks) and adult (9 wks) inbred *Csf1rko* rats exhibited a 3-5 fold increase in circulating granulocytes **(Figure 11B, C**). The *Csf1rko* on the outbred background was reported to be around 70% monocyte-deficient [13]. Monocyte sub-populations were not analysed. In the rat, monocyte sub-populations are distinguished by reciprocal expression of the markers CD43 and HIS48, with HIS48^Hi^ and CD43^Hi^ monocytes corresponding to so-called classical and non-classical monocytes, respectively [32]. Consistent with the role of CSF1R signals in their differentiation, the CD43^Hi^ monocytes were selectively lost in juvenile and adult *Csf1rko* rats. However, there was no corresponding increase in the classical HIS48^Hi^ monocytes (**Figure 11D**). In keeping with previous analysis, there was also a small but significant reduction in circulating B cells whereas T cells were unaffected (**Figure 11E**).

Because of the osteopetrosis, BM cells could only be obtained from *Csf1rko* by crushing the femurs. There was a ∼2-fold reduction in CD45^+^ leukocytes (as a proportion of live cells) recovered from *Csf1rko* BM and a relative increase in granulocytes **(Figure 11F, G**). After excluding granulocytes (HIS48^Hi^/SSC^Hi^), three monocyte/macrophage populations could be distinguished: two putative monocyte populations paralleling peripheral blood HIS48^Hi^ and CD43^Hi^ monocyte profiles, and a CD172A^Low^/CD4^+^ BM resident macrophage population. The classical (HIS48^Hi^) monocytes were more abundant in WT BM than non-classical (CD43^Hi^) monocytes (the reverse of peripheral blood) and their relative abundance was unaffected by the *Csf1rko*. The CD172A^Low^/CD4^+^ resident BM macrophages, as well as B cells, were selectively depleted in *Csf1rko* BM **(Figure 11H, I**).

Surprisingly, the BM transfer did not restore the CD43^Hi^ blood monocyte population in the rescued *Csf1rko* rats whereas the granulocytosis and B cell deficiency in peripheral blood was completely resolved (**Figure 12A-F**). The BM cavity was patent and the populations of IBA1^+^ hematopoietic island macrophages and TRAP^+^ osteoclasts were indistinguishable from WT litter mates (**Figure 12G)**. Accordingly, the yield of CD45^+^ leukocytes from BM obtained by flushing was similar to WT. The CD172^Lo^/CD4^+^/HIS48^-^/CD43^Lo^ resident macrophage population was partially restored and granulocyte and B cell numbers were normalised as in the blood (**Figure 12H-J)**. In standard liquid cultures used routinely to generate BM-derived macrophages in our laboratory [39] WT control BM cells produced a confluent macrophage culture (**Figure 12K**). By contrast, cultures of cells harvested from *Csf1rko* BMT recipients contained only small numbers of large adherent macrophages and there was no increase in cellularity with time.

**Figure 12.**
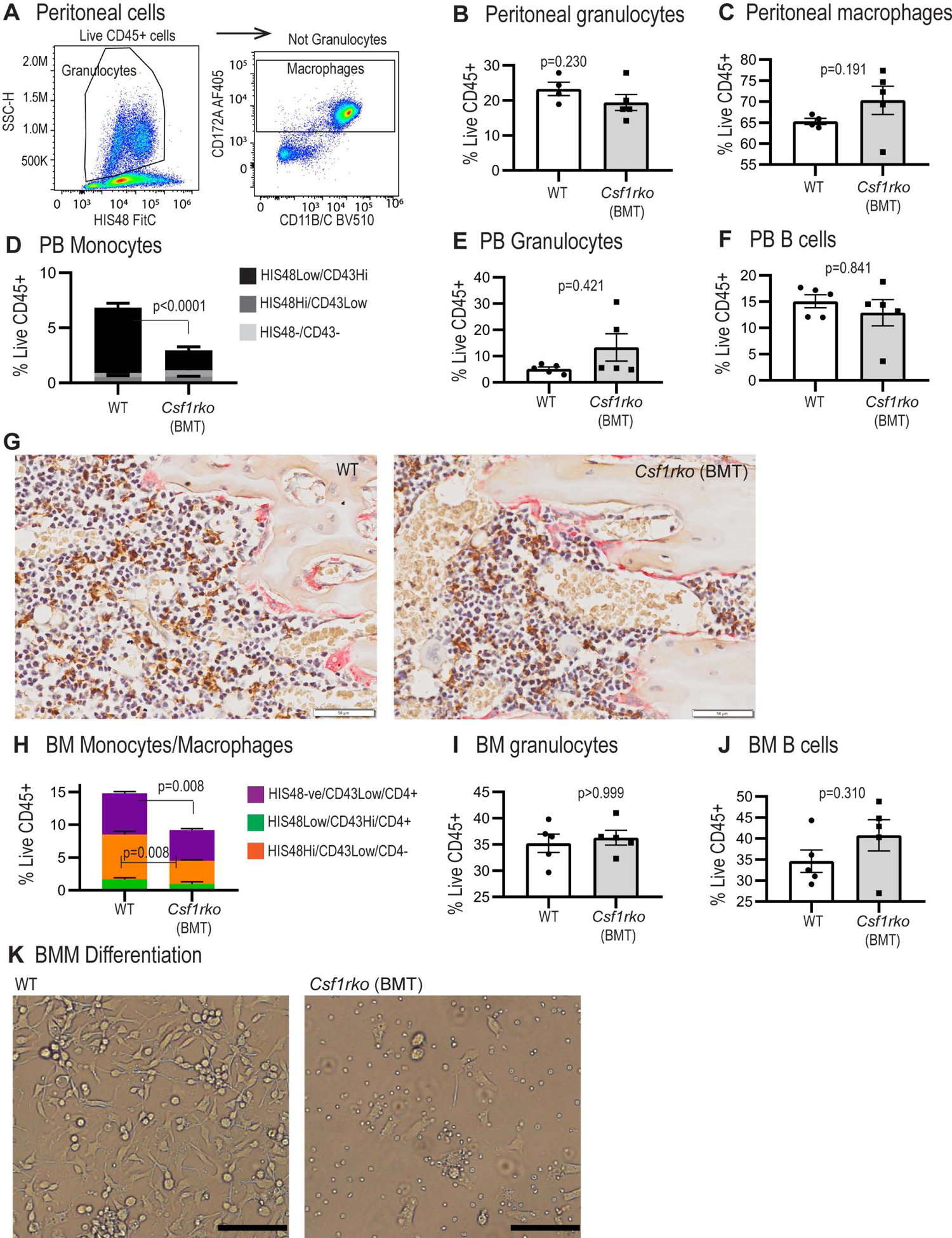
Analysis of the effect of bone marrow transfer at 3 wks of age on peritoneal, peripheral blood and bone marrow leukocyte populations in *Csf1rko* rats. (A) Flow cytometry gating strategy to identify peritoneal granulocytes and macrophages. (B, C) Flow cytometric analysis of (B) peritoneal granulocytes and (C) macrophages in *Csf1rko* rats 9 wks post bone marrow transfer at 3 wks of age compared to age-matched WT controls. (D-F) Flow cytometric analysis of peripheral blood (D) monocyte subsets, (E) granulocytes and (F) B cells in *Csf1rko* rats 9 wks post bone marrow transfer at 3 wks of age compared to age-matched WT controls. (G) Representative images showing dual staining for IBA1 expression (brown) and TRAP activity (magenta) in the tibiae of *Csf1rko* rats at 6-weeks post-BMT (right) and aged matched WT control (9 weeks) (left). Original magnification: 40X. Scale bar: 50µm. (H-J) Flow cytometric analysis of bone marrow (BM) (G) monocyte/macrophage subsets, (H) granulocytes and (I) B cells in *Csf1rko* rats 9 wks post bone marrow transfer at 3 wks of age compared to age-matched WT controls. (K) BM from WT and *Csf1rko* rats shown in panels G-H was cultured for 7 days in 100 ng/ml CSF1 to generate bone marrow-derived macrophages. Images shown are representative of cultures from 5 separate WT and BMT recipient *Csf1rko* animals (scale bar 50 µM). All genotype comparisons were analysed by Mann Whitney test.

## Discussion

### The role of CSF1R in postnatal growth

Our results demonstrate that CSF1R-dependent macrophages are essential for early postnatal growth and organ development in the rat. The exclusive impact on postnatal growth distinguishes the *Csf1rko* from *Igf1* and *Igf2* mutations which impact the growth of the embryo [40]. Interestingly, although *Csf1r* is highly-expressed in placenta, the lack of impact of the mutation on embryonic growth also indicates that placental function is CSF1R-independent. It is likely that any loss of macrophage-derived trophic factors in the embryo is mitigated to some extent by placental and maternal-derived growth factors. Severe postnatal growth retardation is not evident in human patients with bi-allelic *CSF1R* mutations [8, 10, 11, 19]. This may be indirect evidence that the mutant alleles in these individuals are hypomorphic, rather than complete loss-of-function. The only definitive human homozygous *CSF1R* null mutation described thus far was associated with severe osteopetrosis, brain developmental defects and infant mortality [41].

The somatomedin hypothesis proposed that somatic growth is controlled by pituitary GH acting on the liver to control the production of IGF1. Numerous analyses of conditional mutations of *Ghr, Igf1* and *Igf1r* in various tissues and cell types in mice paint a more complex picture [26, 42, 43]. Notably, conditional deletions of *Igf1* and *Igf1r* in chondrocyte and osteoblast lineage cells produce substantial reductions in overall somatic growth rates (reviewed in [44]). Hence, as suggested by Chitu & Stanley [3], the defects in skeletal growth and maturation that we showed were already evident in the *Csf1rko* at birth (**Figure 1)** are likely one underlying cause of reduced postnatal somatic growth rate.

By contrast, conditional deletion of *Igf1* in the mouse liver had a marginal effect on somatic growth despite 70-90% loss of circulating IGF1 [42, 43]. Like the GH-deficient dwarf rat [45] and GHR-deficient mice [46] the *Csf1rko* rats have greatly-reduced circulating IGF1. The loss of IGF1 was also present, albeit less severe, on the outbred background. However, the *Csf1rko* rats are not GH-deficient, consistent with unchanged levels of GH mRNA in the pituitary [13, 18], and the phenotypes of GH and *Csf1rko* mutations are quite distinct. Dwarf rats have a slower overall growth rate but do not exhibit the early growth arrest and major organ phenotypes and morbidity seen in the *Csf1rko* irrespective of genetic background. Indeed, GH/GHR deficiency in rats and mice is associated with increased longevity [47]. Macrophages themselves produce an array of growth factors, including IGF1, that are implicated in development and tissue repair [7, 48]. Meta-analysis of mouse resident tissue macrophage gene expression [33] revealed constitutive high expression of *Igf1* and *Igfbp4* mRNA but macrophages are clearly not the only extrahepatic source. Rat macrophages grown in CSF1 also express abundant *Igf1* mRNA [39] and given their relative abundance, could be a significant source of IGF1 in tissues. However, conditional deletion of *Igf1* in myeloid cells in mice had no reported effect on somatic growth [49]. Furthermore, depletion of microglia in the brain of the *Csf1rko* rat had no effect on *Igf1* mRNA [13, 18]. It is notable that the phenotypes we have described in the *Csf1rko* rat are considerably more severe than those reported in *Csf1*^tl/tl^ rats, which are also deficient in circulating IGF1 [34]. As described originally [50] and confirmed on an inbred background [51], *Csf1*^tl/tl^ rats achieve body weights of 250-300g, are male fertile and have normal longevity. This suggests that in rats the alternative ligand, IL34, has significant non-redundant roles in macrophage homeostasis and function in multiple organs during postnatal development. Accordingly, we suggest that CSF1R-dependent macrophages act indirectly to regulate circulating IGF1 mainly through their effects on hepatocyte proliferation and maturation but this is not linked directly to the impacts on somatic growth.

### The effect of the *Csf1rko* on the liver

The set of hepatic genes correlated with *Csf1r* in network analysis (**Figure 4A, Table S1C)** includes *Cd68* and *Aif1,* encoding CD68 and IBA1 respectively, and confirms a rat Kupffer cell (KC) signature [22] that includes marker genes enriched in this resident population in mice (e.g. *Cd5l, Clec4f, Timd4, Vsig4*) [52, 53] as well as highly-expressed genes involved in iron metabolism/erythrophagocytosis (*Cd163, Slc40a1, Timd2*). The liver has a unique ability to regenerate following partial hepatectomy to return to a constant liver-body weight ratio, a so-called hepatostat [20]. Administration of CSF1 to adult mice, rats or pigs [32, 54–56] overcomes this constraint, drives hepatocyte proliferation and accelerates regeneration following partial hepatectomy.

During the postnatal period in rat liver a phase of hyperplasia is followed by hypertrophy and structural maturation to form mature hepatic sinusoids lined by a single layer of hepatocytes [23]. In addition to impacts of the *Csf1rko* on the IGF1 axis in the liver, transcriptional profiling indicated a broader delay in this postnatal hepatocyte maturation exemplified by the retention of fetal liver-expressed genes such as lipoprotein lipase (*Lpl*)[57], fetal liver hepcidin (*Hamp*) [58] and the fetal amino acid transporter *Slc38a2* [59]. Two other genes highly enriched in the liver of *Csf1rko* rats were the cold shock-inducible gene *Rbm3* and stress-inducible *Gadd45b.* Expression of *Rbm3* and *Gadd45b* and their potential impacts on liver metabolism [60, 61] could be a consequence and/or a cause of the reduced adiposity and hepatic steatosis of the *Csf1rko.* Conversely, the relative loss of several highly expressed liver-specific genes is likely to exert additional pleiotropic effects on other organs. Amongst the most *Csf1r-*dependent transcripts, SERPINA6 (transcortin) is the major binding protein for circulating glucocorticoids implicated in regulation of the response to GH [62]. In mice, *Serpina6* is highly-expressed in fetal liver, declines to almost undetectable level at birth, and is then re-induced in parallel with *Ghr* and *Igf1* [25].

The liver of 7-day old rats contains abundant lipid droplets [27] but this usually resolves rapidly. The coordinated reduction of numerous genes involved in lipid metabolism (**Figure 4A)** in the liver in 3 wks old *Csf1rko* rats is the reciprocal of the increase observed following CSF1 treatment of neonates [27] and likely contributes to both the progressive steatosis and the lack of visceral adipose in the animals. Both of these phenotypes were reversed by the transfer of wild type BM cells (**Figure 7 and 10**). The coordinated regulation of genes involved in lipid metabolism may be related to the reduced expression of the transcriptional repressor *Hes6* [63] in the *Csf1rko*.

The precise mechanisms underlying reduced proliferation in the postnatal *Csf1rko* liver are not evident from the transcriptome analysis, which did not reveal the loss of known growth regulators or indirect evidence of deficient receptor signalling. Expression of potential candidates is highlighted separately in **Table S1E**. It is also notable that the *Csf1rko* has no impact on expression of markers of other non-parenchymal cells, endothelial cells (*Pecam1, Cdh5*) or hepatic stellate cells (*Pdgfrb*). Postnatal growth of the liver involves β−catenin signalling linked to E-cadherin and the receptor for hepatocyte growth factor (MET). Conditional deletion of *Ctnnb1* in hepatocytes led to a 15-25% decrease in LBW ratio at postnatal day 30 [64]. Conditional deletion of *Yap1,* a downstream component of the hippo kinase pathway also led to reduced hepatocyte proliferation and reduced LBW [24]. By contrast, the grossly-reduced hepatocyte proliferation seen in the juvenile *Csf1rko* is not liver-specific and does not lead to reduced LBW ratio. The regulation of hepatocyte proliferation has been studied mainly in the context of regeneration and involves a complex array of growth factors; both locally-derived and present in the circulation (reviewed in [21]). As noted in the introduction, CSF1 treatment of newborn rats can promote the selective growth of the liver [27]. However, the intrinsic hepatostat remains functional in the *Csf1rko* rat and the reduced hepatocyte proliferation may partly be a consequence of deficient somatic growth that is also CSF1R-dependent.

### The effect of Csf1rko on skeletal development

Although there are similarities in some phenotypes, the osteosclerotic bone phenotype in the *Csf1rko* is quite distinct from osteopenia associated with GH or IGF1 deficiency [65]. The novel phenotype we identified in the cranial case relates to the process of intramembranous ossification. Intramembranous bones ossify directly from preosteogenic condensations of multipotent mesenchymal stem cells without a chondrocyte intermediate. Like endochondral ossification, this process is dependent upon angiogenesis (reviewed in [66]). The process of bone formation in the cranial case is initiated by postnatal expansion of bone from the sutures formed during embryonic development [67]. Osteoblast maintenance and calcification requires input from osteomacs, a bone-associated macrophage population distinct from osteoclasts [68]. These CSF1-responsive macrophages promote the processes of both endochondral and intramembranous ossification during bone repair *in vivo* [69, 70]. Analysis of the *Csf1*^tl/tl^ rat revealed a defect in osteoblast attachment to bone surfaces and the absence of prominent stress fibres[71]. Hence, the selective failure of intramembranous ossification in the cranial case, and the increased calcification of the base of the skull, may reflect distinct functions of osteomacs and osteoclasts, both dependent on CSF1R.

The *Csf1rko* rat resembles bi-allelic human *CSF1R* mutation in the pronounced under-modelling of the digits [8]. Detailed analysis of digital development in the mouse [72] revealed morphologic and calcification patterns in the subarticular regions that were also distinct from archetypal physeal endochondral ossification. The defects we observe in the *Csf1rko* rats resemble skeletal dysplasia caused by mutations in the gene encoding the macrophage-enriched transcription factor MAFB [72] and more generally various forms of malignant osteopetrosis (reviewed in [73]).

### The role of CSF1R-dependent macrophages in development of other organ systems

The combined analysis using the *Csf1r-*mApple reporter (**Figure 6)** and localization of IBA1 (**Figures 8, 9**) highlights both the abundance of resident tissue mononuclear phagocytes in WT rats and the almost complete depletion of these cells in the *Csf1rko* regardless of genetic background. The loss of monocytes in *Csf1rko* rats [13] was shown here to be selective for the CD43^Hi^ ‘non-classical’ monocyte population which is the major population in the rat [32]. Analysis of a hypomorphic *Csf1r* mutation [36] and the selective impact of anti-CSF1R antibody [74] indicates that CSF1R is not required for monocyte production, whereas tissue macrophages require CSF1R signals for survival. The lack of accumulation of classical (His48^Hi^) monocytes in the *Csf1rko* despite the block to maturation could reflect more rapid transit of these cells in the circulation. For example, the striking periportal concentration of the residual IBA1^+^ Kupffer cells in the liver and reduced numbers in the gut lamina propria could be associated with continuous extravasation of monocytes and subsequent short half-life in the tissue.

In many different organ systems, depletion of resident tissue macrophages permits monocyte extravasation to occupy the vacant territory or niche (reviewed in [1, 2]). It is rather striking that the rescue of tissue macrophage populations following BMT (**Figure 10**) restores them to almost precisely wild-type levels. Guilliams *et al.* [1] favour the idea that each macrophage occupies a defined niche responding mainly to local CSF1. An alternative view is that the macrophages are territorial and the regular distribution in every tissue (confirmed in **Figures 8-10**) is determined by mutual repulsion [2]. That view favours a role for systemic CSF1 stimulation. Based upon the rapid reduction in circulating CSF1 in the BMT recipients **(Figure S5A)** and the ability of exogenous CSF1 to expand macrophage populations in all tissues [32] we suggest that the entire mononuclear phagocyte system is regulated in a coordinated manner by CSF1 availability. Following BMT, macrophage repopulation occurs until balance is restored between CSF1 production and CSF1R-mediated clearance/utilisation.

Analysis of the role of macrophages in postnatal development based upon *Csf1r* and *Csf1* mutations in mice has focussed on one organ at a time, emphasising local functions [3]. The loss of circulating IGF1 is just one example of the systemic consequences of the *Csf1rko.* We clearly cannot separate direct local roles of macrophages in the regulation of proliferation and differentiation in liver, brain, skeleton, muscle, lung, kidney, gut, thymus, adipose and gonads from systemic consequences of developmental abnormalities in every other organ. For example, although growth hormone (*Gh*) mRNA was not reduced in the pituitary of *Csf1rko* rats, RNA-seq profiling revealed a significant reduction in cell-cycle associated transcripts and relative reductions in *Fshb* and *Lh* in males and prolactin (*Prl*) in both sexes [18].

Given the large numbers of macrophages in the developing embryo and their roles in clearance of apoptotic cells [16] it is surprising that there is so little apparent impact of their almost complete absence in the *Csf1rko* on prenatal development. “Amateur” phagocytes can evidently replace the phagocytic functions of macrophages in embryonic development. In keeping with this view, although genes encoding lysosomal enzymes and endosome-associated proteins are highly-expressed by microglia, the relative expression of these genes was unaffected in brains of *Csf1rko* rats or microglia-deficient mice [13, 36].

### The nature of the progenitor cells that mediate phenotypic rescue

Bennett *et al* [14] reported that pre-weaning lethality could be prevented in around 50% of inbred *Csf1rko* mice by neonatal transfer of WT BM cells. Their study focussed on reconstitution of microglia and did not examine the contribution of the donor cells to the hematopoietic compartment. Importantly, they showed that rescue was independent of the monocyte chemotactic receptor, CCR2. Like these authors, we were able to fully repopulate a myeloid population in the brain following BMT in a *Csf1rko* recipient (**Figure 10B**). The less-ramified stellate morphology we observed was not reported directly in the mouse study, but is evident in the published images [14]. Ongoing studies address the question of whether these BM-derived “microglia” reconstitute expression of the microglial gene expression signature that is lost in the *Csf1rko* brain [18].

In principle, neonatal transfer in the published study [14] could allow WT hematopoietic stem cells (HSC) to populate the developing bone marrow. In the rat, we were actually able to reverse and rescue the *Csf1rko* phenotype and resident tissue macrophage populations, including those of the brain with 100% success by transfer at weaning when the developmental defects including severe osteopetrosis were already evident. The recipients were not treated to deplete HSC and were not deficient in other blood cell lineages. The reversal of osteosclerosis and expansion of the hematopoietic niche in the long bones likely requires the generation of CSF1R-dependent osteoclasts and resident osteomacs [38, 75] that were replenished in the BMT recipients (**Figure 12**). Similarly, CSF1 treatment restored TRAP^+^ osteoclast populations and active resorption in the bone of *Csf1*^tl/tl^ rats [76]. The granulocytosis and B cell deficits observed in marrow and blood of the *Csf1rko* rats were reversed by BMT despite the lack of contribution of donor cells to the BM compartment. We eliminated any increase in the key growth factor, CSF3, as a mechanism underlying granulocytosis (**Figure S5**). Neutrophils have a short half-life in the circulation [77]. In part, the granulocytosis in the *Csf1rko* rat probably reflects the loss of Kupffer cells and selected splenic macrophages which clear senescent neutrophils via phosphatidylserine-mediated phagocytosis [78]. The *Csf1r-*mApple reporter transgene is expressed by neutrophils, but in most tissues of the *Csf1rko* rat there was no evidence of any residual positive myeloid cells (**Figure 6**) suggesting that the loss of macrophages also compromised neutrophil extravasation. Neutrophils migrate constitutively into healthy tissues and their homeostasis depends upon the CXCR4/CXCL12 axis [77]. Interestingly, *Cxcl12* mRNA was highly-expressed by rat liver and down-regulated in the *Csf1rko* (**Table S1**).

The lack of CSF1 responsiveness in BM in the rescued *Csf1rko* rats (**Figure 12K**) is consistent with the failure of the BMT to rescue circulating monocyte numbers and indicates that the transplant does not contribute to the HSC and myeloid progenitor populations. This is not surprising since the recipients received no conditioning and there is no evidence of hematopoietic insufficiency. The implication is that there is a committed progenitor in rat BM that can directly provide long term engraftment of tissue mononuclear phagocyte populations without a monocyte intermediate. The ability of hematopoietic stem and progenitor cells to traffic through lymph and blood and to populate resident myeloid populations directly has been reported in mice [79]. The key donor population may be related to the CSF1-responsive clonogenic monocyte/macrophage progenitor described in mouse BM [80]. On the other hand, a recent study indicated that descendants of the yolk sac erythro-myeloid progenitor that can give rise to tissue macrophages and osteoclasts may be present in the circulation [81]. Studies in the chick also revealed the existence of a BM progenitor that could produce long term macrophage-restricted chimerism when injected into the embryo prior to the onset of definitive hematopoiesis [4]. Further characterization of BM macrophage progenitor populations in the rat will depend on development of markers that are not currently available. The current study demonstrates the potential utility of the *Csf1r-*mApple transgene, which is currently being backcrossed to the DA background.

The second key implication of the rescue of the *Csf1rko* relates to the origins of resident macrophages. There is an emerging view based upon lineage trace models in mice that most tissue macrophages are seeded during embryonic development and thereafter maintained by self-renewal [1, 2]. Whether or not this model can be extended to other species, the long-lived effectiveness of the rescue of *Csf1rko* rats indicates that macrophage territories in every organ can be occupied by cells of BM origin and maintained in the absence of CSF1R-expressing monocytes.

In conclusion, we have shown that severe developmental abnormalities in inbred *Csf1rko* rats can be reversed by WT bone marrow. The phenotypic rescue achieved by BMT is consistent with evidence that CSF1R expression is entirely restricted to MPS lineage cells (Reviewed in [6, 82] and all impacts of *Csf1rko* mutation are attributable to their absence.

## Material and Methods

### Generation of transgenic rats and animal maintenance

To create a pure DA line the original outbred *Csf1rko* was backcrossed to WT DA (Animal Resource Centre, Perth, Australia) for at least 7 generations. As *Csf1rko* rats do not develop teeth, to ensure their survival and maximise their growth, we routinely separate them from their littermates at 3 wks and commence a feeding regime including wet mashed standard chow and a veterinary powdered milk nutritional supplement (Di-Vetelact, Sydney, Australia). The same approach is used in maintenance of *Csf1^op/op^* mice (e.g. [28]). Rats were bred and maintained in specific pathogen free facilities at The University of Queensland (UQ) under protocols approved by the UQ Animal Ethics Unit.

### Bone marrow transfer

Bone marrow was obtained from the femurs and tibias of WT rats by flushing with a 26G needle. 1×10^7^ bone marrow cells in saline solution containing 2% fetal bovine Serum (FBS) were transferred to 3 wk-old *Csf1rko* recipients by intraperitoneal injection. Recipients were maintained on the supplemented diet.

### Flow cytometry

100 ul peripheral blood was collected into ethylenediaminetetraacetic acid (EDTA) tubes by cardiac puncture after CO_2_ euthanasia. Red cells were lysed for 2 min in ACK lysis buffer (150mM NH_4_Cl, 10mM KHCO_3_, 0.1mM EDTA, pH7.4), the leukocytes were centrifuged and washed twice in PBS and the pellet resuspended in flow cytometry (FC) buffer (PBS/2%FBS) for staining. Cells were stained for 45 min on ice in FC buffer containing unlabelled CD32 (BD Biosciences, Sydney, Australia) to block Fc receptor binding and HIS48-FitC, CD11B/C-BV510, CD45R-PE (BD Biosciences), CD43-AF647, CD4-APC/Cy7 (Biolegend, San Diego, CA, USA) and CD172A-AF405 (Novus, Noble Park North, Australia). Cells were washed twice and resuspended in FC buffer containing 7AAD (LifeTechnologies, Musgrave, Australia) for acquisition on a Cytoflex flow cytometer (Beckman Coulter Life Sciences, Sydney, Australia). Single colour controls were used for compensation and fluorescence-minus-one controls were used to confirm gating. Data were analysed using FlowJo 10 (Tree Star; https://www.flowjo.com).

### Micro-CT imaging and reconstruction

Bones for micro-CT were fixed in 4% paraformaldehyde (PFA) and scanned using Bruker’s Skyscan 1272 (Bruker, Belgium) by rotating over 360° in 0.8° rotational steps. The X-ray settings were standardised to 70 kV and 142 µA with an exposure time of 1450 ms and the X-ray filter used was a 0.5 mm aluminium. Projections were acquired with nominal resolutions of 21.5 and each resulting image contained 1144 x 1144 pixels. All X-ray projections were reconstructed using NRecon 1.7.3.1 software (Bruker) to create cross-sectional images and viewed using CTvox 3.3.0 (Bruker).

### Bone immunohistochemistry (IHC) and histological staining

Bone tissues were processed as per previously [70]. Fixed bones were decalcified in 14% EDTA for 8-10 wks. IHC was performed on deparaffinized and rehydrated 5 µm sections that were blocked for endogenous peroxidase activity using 3% hydrogen peroxide. Antigens were retrieved using 0.1% trypsin (Sigma-Aldrich, MO, USA) and non-specific staining was blocked using 10% fetal bovine serum/neonatal goat serum for 1 hr prior 90 min incubation with either biotinylated anti-laminin antibody (Novus Biologicals, Colorado, USA) or unconjugated primary antibody against IBA1 (FUJIFILM Wako Chemicals, VA, USA). For IBA1 staining sections were subsequently incubated with a biotinylated F(ab’)2-Goat anti-Rabbit IgG (Vector Labs, CA, USA). All sections were then incubated with horseradish peroxidase (HRP)-conjugated streptavidin (Dako Agilent Pathology Solutions, DK). Diaminobenzidine was developed as per the manufacturer’s instructions (Vector Labs) and all sections were counterstained with Mayer’s hematoxylin (Sigma-Aldrich).

Tartrate-resistant acid phosphatase (TRAP) activity was detected as previously described [83]. For dual IBA1 and TRAP staining, IBA1 staining was performed before TRAP detection. Whole-mount Alcian Blue-Alizarin Red staining of newborn rats was performed as previously described [84]. Staining was imaged using VS120 slide scanner (Olympus, Tokyo, Japan) or SZX10 stereo microscope with DP26 digital camera (Olympus). Laminin-stained sections from 3-week-old rat limbs were used to examine the muscle fiber diameter of the posterior tibialis. The average diameter of 30 muscle fibers was analyzed using Visiopharm® software (Visiopharm, Hørsholm, Denmark).

### Brain immunohistochemistry

Rat brains were harvested and fixed in 4% paraformaldehyde for 48 h and then transferred into PBS containing 0.01% azide. Brains were sectioned in the sagittal plane using a vibratome (Leica VT 1200S, Leica Biosystems, Mt Waverley, Australia). Free-floating sections were first incubated at room temperature for 30 min in permeabilization buffer [1 % Triton and 0.1 % Tween20 in PBS] followed by 30 min in blocking solution [4 % FCS,0.3 % Triton and 0.05 % Tween20 in PBS]. Sections were then incubated for 24 h at room temperature under orbital agitation in either rabbit anti NeuN (Millipore ab10807945, Melbourne, Australia, Lot# ABN78, 1:500) or rabbit anti IBA-1 (Wako AB_2314666, USA, Cat# 01-1874, 1:500) diluted in blocking solution. After 3 × 10 min washes in blocking solution, slices were incubated in goat anti-rabbit-Alexa 488 (Thermofisher Scientific, Brisbane, Australia, 1:1000); diluted in blocking solution for 4 h at room temperature. Slices were then washed in PBS (3 × 10 min) followed by a 5 min incubation with DAPI (Thermofisher Scientific, 1: 5000) diluted in PBS. All sections were washed once with PBS for 5 min and mounted with Fluorescence Mounting Medium (Dako, Agilent, Santa Clara, California, USA). Images were acquired on a fluorescent slide scanner (Zeiss Axioscan, Zeiss, Sydney, Australia) with either a 10X or 40X objective (1024 × 1024).

### Immunohistochemistry of other tissues

Tissues were harvested and fixed in 4% paraformaldehyde for 24 h and then processed for paraffin-embedded histology using routine methods. Sections were deparaffinised and rehydrated in descending ethanol series. For H&E, 4 µm sections were stained with eosin and hematoxylin (Sigma-Aldrich) for 30 second and 1 minute respectively. For elastin staining, 4 µm sections were stained with Elastin solution, Weigert’s iron hematoxylin solution and Picrofuchsin solution for 10, 5 and 2 minutes respectively (Elastin van Gieson staining kit, Merck, Melbourne, Australia). For Ki67 and IBA1 staining, epitope retrieval was performed by heat induction in Diva Decloaker (DV2004MX, Lot:011519, Biocare Medical, California, USA). Sections were stained with rabbit anti Ki67 (Abcam ab16667, Cambridge, UK, Lot# GR331319528, 1:100) or rabbit anti-IBA1L (FUJIFILM, Wako Chemicals, Richmond, Virginia, USA, 019-19741, Lot# CAK1997, 1:1000). Secondary detection was with DAKO Envision anti rabbit HRP detection reagents (Agilent).

Sections were then dehydrated in ascending ethanol series, clarified with xylene and mounted with DPX mountant (Sigma-Aldrich). Whole-slide digital imaging was performed on the VS120 Olympus slide scanner. DAB-positive areas were quantified using ImageJ (https://imagej.net/) in six different field per sample. For lung, DAB-positive areas were normalised to total number of cells per field.

### RNA purification and qRT-PCR analysis

RNA was extracted using TRI Reagent (Sigma-Aldrich), according to manufacturer’s instructions. For each extraction ∼100 mg of tissue was processed using 1 ml reagent. Genomic DNA contamination of RNA preparations was eliminated by digestion with DNase I amplification grade (Thermofisher Scientific). RNA quantity was measured using a Nanodrop and RNA integrity estimated on an Agilent ≥2200 Tapestation System (Agilent). The RNA Integrity Number (RINe) was calculated for each sample and all samples used for sequencing had a RINe of ≥ 7.0.

Expression levels for selected genes were quantified using real-time PCR. cDNA was synthesized from 1 µg total RNA using cDNA synthesis kit (Bioline, Sydney, Australia) and RT-PCR was performed using the SYBR® Select Master Mix (Thermofisher Scientific) on an Applied Biosystems™ QuantStudio™ real-time PCR system (Thermofisher Scientific). Gene expression relative to *Hprt* was calculated using the ΔCt method. Primer sequences: *Hprt* F: CTCAGTCCCAGCGTCGTGA, R: AACACCTTTTCCAAATCTTCAGCA; *Adgre1* F: GGGGCTATGGAATGCATAATCGC, R: AAGGAGGGCAGAGTTGATCGTG; *Csf1r* F: GACTGGAGAGGAGAGAGCAGGAC, R: CTGCCACCACCACTGTCACT.

### Library preparation and sequencing

RNA-seq libraries were prepared by IMB Sequencing Facility, University of Queensland, Australia, with the TruSeq Stranded mRNA Library protocol (Illumina, San Diego, California, USA). Sequencing of 12 liver samples (with 84 unrelated samples) was performed using a single NovaSeq S1 200 cycle sequencing run on a NovaSeq 6000 machine (Illumina) through IMB Sequencing Facility. Sequencing depth was between 9 million and 35 million paired end reads per sample. The raw sequencing data, in the form of .fastq files, are deposited in the European Nucleotide Archive under study accession number PRJEB39130.

### Read pre-processing

Reads were pre-processed using fastp v0.20.0 [85] with parameters --length_required 50 -- average_qual 10 --low_complexity_filter --correction --cut_right --cut_tail --cut_tail_window_size 1 --cut_tail_mean_quality 20. These parameters: (1) trim all bases from the 3’ end that have quality < 20; (2) cut reads should the mean quality within a 4bp window, advanced 5’ to 3’, fall below 20; (3) require that 30% or more of the bases in each read are followed by a different base (an indicator of read complexity); (4) require a mean base quality across the entire read of > 10. By default, fastp also trims auto-detected adapters, discards reads with > 5 N (undetermined) bases, and requires that > 40% of the bases in each read have Phred score >15. Mismatched base pairs were corrected in regions of paired end reads that overlapped each other, should one base have a quality score higher than the other. This required a minimum overlap of 30bp, a difference in base qualities of 5, and no more than 20% of the bases in the overlapping region needing correction. Finally, we required a minimum read length of 50bp. Pre-processing discarded on average 8.5% of the bases per sample (Supplementary Table 1).

### Expression quantification and differential gene expression analysis

For each set of cleaned reads, expression level was quantified as transcripts per million (TPM) using Kallisto v0.44.0 [86] with 100 bootstrap replicates. Kallisto quantifies expression by reference to an index of transcripts, which was constructed using the combined set of unique protein-coding transcripts from the Ensembl (ftp://ftp.ensembl.org/pub/release-98/fasta/rattus_norvegicus/cdna/Rattus_norvegicus.Rnor_6.0.cdna.all.fa.gz) and NCBI RefSeq (ftp://ftp.ncbi.nlm.nih.gov/genomes/all/GCF/000/001/895/GCF_000001895.5_Rnor_6.0/GCF_0000 01895.5_Rnor_6.0_rna.fna.gz) versions of the Rnor_6.0 annotation, as previously described [39] (n = 25,870 protein-coding genes in total, of which 22,238 had Ensembl IDs).

Differential expression of genes comparing WT and *Csf1rko* was performed using Degust (https://degust.erc.monash.edu/). The FDR cut-off was 0.05, the method was Voom/Limma and only samples with a minimum read count of 1 TPM in at least one replicate were included.

### Network analysis of gene expression

Network cluster analysis of gene expression in the livers of *Csf1rko* and WT rats was performed using BioLayout (http://biolayout.org). The expression levels determined by Kallisto were filtered to remove any gene where no sample reached 1 TPM. Similarities between expression profiles of individual samples (sample to sample analysis) or genes (gene to gene analysis) were determined by calculating a Pearson correlation matrix. For the sample to sample analysis, relationships where r ≥ 0.93 (the highest r which included all 12 samples) were included. For the gene to gene analysis, the results were filtered to remove all relationships where r < 0.95. A network graph was constructed by connecting the remaining nodes (samples or genes) with edges (where the correlation coefficient between two samples or genes exceeded the threshold value). Samples or genes with similar expression patterns were located close to each other in the network. The gene to gene network graph was interpreted using the Markov Cluster Algorithm (MCL) at an inflation value (which determines cluster granularity) of 1.7. Genes with similar expression patterns were allocated to the same cluster, forming cliques of interconnected nodes. All results were verified using Spearman (non-parametric) correlation coefficient.

### IGF1, GH, CSF1 and CSF3 Immunoassay

Serum IGF1 was measured using Mouse/Rat DuoSet ELISA kit (R&D Systems, Minneapolis, Minnesota, USA) according to manufacturer instruction. GH was measured in rat serum as described previously [87]. Briefly, an ELISA plate was coated overnight at 4°C with capture antibody (National Institute of Diabetes and Digestive and Kidney Diseases (NIDDK)-anti-rat GH (rGH)-IC-1 (monkey), AFP411S, NIDDK-National Hormone and Pituitary Program (NHPP, Torrance, California, USA) at a final dilution of 1:40,000. Non-specific binding was blocked using 5% skim milk in 0.05% PBS with Tween-20 for 2 hours. The bound protein was detected using rabbit anti-GH antibody at a final dilution of 1:40,000 (AFP5672099, NIDDK-NHPP) followed by horseradish peroxidase-conjugated antibody (anti-rabbit, IgG; GE Healthcare, Chicago, Illinois, USA) at a final dilution of 1:2000. The concentration of growth hormone was calculated by regression of the standard curve generated using a 2-fold serial dilution of mouse GH (mGH) (AFP-10783B, NIDDK-NHPP) in PBS-T supplemented with 0.2% BSA. Serum CSF1 and CSF3 were analysed using Rat CSF1 SimpleStep ELISA kit (Abcam, Australia, Cat# ab253214) and Rat Granulocyte Colony Stimulating Factor (G-CSF/CSF3) ELISA Kit (MyBioSource, San Diego, CA, USA, Cat# MBS265555) respectively according to manufacturer instructions.

### Glucose and Insulin measurements

Blood glucose and serum insulin were measured in non-fasted animals and in accordance with the manufacturer’s instructions. Blood glucose was measured using a Sensocard glucometer and glucose strips (Point of Care Diagnostics, Sydney, Australia). Insulin levels were measured using a mouse and rat insulin ELISA (Mercodia, Uppsala, Sweden).

### Quantitative measures of digestive tract

Villi and crypt length and villi width in the small intestines of a mix of male and female *Csf1rko* (n=6) or WT (n=9) rats on the Dark Agouti background were measured as previously described [88]. 100-200 villi and crypts were analysed per rat ileal section and averaged for analysis. For quality control, only villi and crypts where a continuous epithelium from the base of the crypts to the tip of the villi could be measured were included in the analysis. For width analysis, total IBA^+^ immunolabelling was also analysed in villus lamina propria (VLP), and the subepithelial dome (SED) and follicular-associated epithelium (FAE) of Peyer’s patches (PP; 1-4 PP/rat). For submucosa and muscularis analysis, thickness for each was measured at 25 random spots on each rat ileal section.

### TUNEL staining

TUNEL staining was performed using TACS® 2 TdT DAB (diaminobenzidine) Kit (Trevigen, Gaithersburg, Maryland, USA) according to manufacturer instruction.

### Mammary gland wholemount

Mammary gland wholemounts were prepared using CUBIC-based tissue clearing and methyl green staining, as previously described [89]. Glands were imaged using an Olympus SZX10 stereo microscope. Exposure time was consistent between rat samples and brightness and contrast adjustment uniformly applied. Ductal and fat pad length were measured from the #4 mammary glands.

### Mammary tissue immunohistochemistry

Immunostaining on formalin-fixed paraffin embedded rat mammary tissue was performed as previously described [90]. The following primary antibodies were used in this study: chicken anti-KRT5 (Biolegend, San Diego, California, USA; 905903, Lot# B279859, 1:1000), mouse anti-E-cadherin (BD Biosciences; 610182, Lot# 7292525, 1:400) and rabbit IBA1 (FUJIFILM Wako Chemicals; 019-19741, Lot# CAK1997, 1:1000). The following secondary antibodies (Thermofisher Scientific) were used in this study (1:500): goat anti-chicken AF488 (A11039), goat anti-mouse AF555 (A32727) and goat anti-rabbit AF647 (A21245). Nuclei were stained with DAPI dilactate (625 ng/mL) for 10 min. Tissue was imaged using an Olympus BX63F upright epifluorescence microscope. Brightness and contrast was optimally adjusted for lineage markers and DAPI. Exposure, brightness and contrast were kept consistent between tissue sections for IBA1.

## Statistics

Statistical tests were performed using GraphPad Prism 7.03. Comparisons between WT and *Csf1rko* were performed using the unpaired Student’s t test or Mann-Whitney test as indicated in Figure Legends.

## Acknowledgments

We would like to thank Ms. Lisa Foster (Manager, UQ Biological Resources) and her staff (especially Rachel Smith) for assistance with breeding and husbandry of the *Csf1rko* rats. The generation of the *Csf1rko* rat was supported by UK Medical Research Council Grant MR/M019969/1 to DAH and CP. This work was supported in part by Australian National Health and Medical Research Council (NHMRC) Grant GNT1163981 awarded to DAH and KMS. The laboratory is grateful for core support from The Mater Foundation. We acknowledge support from the Microscopy and Cytometry facilities of the Translational Research Institute (TRI). TRI is supported by the Australian Government.

## Supplementary Figures

**Figure S1.**
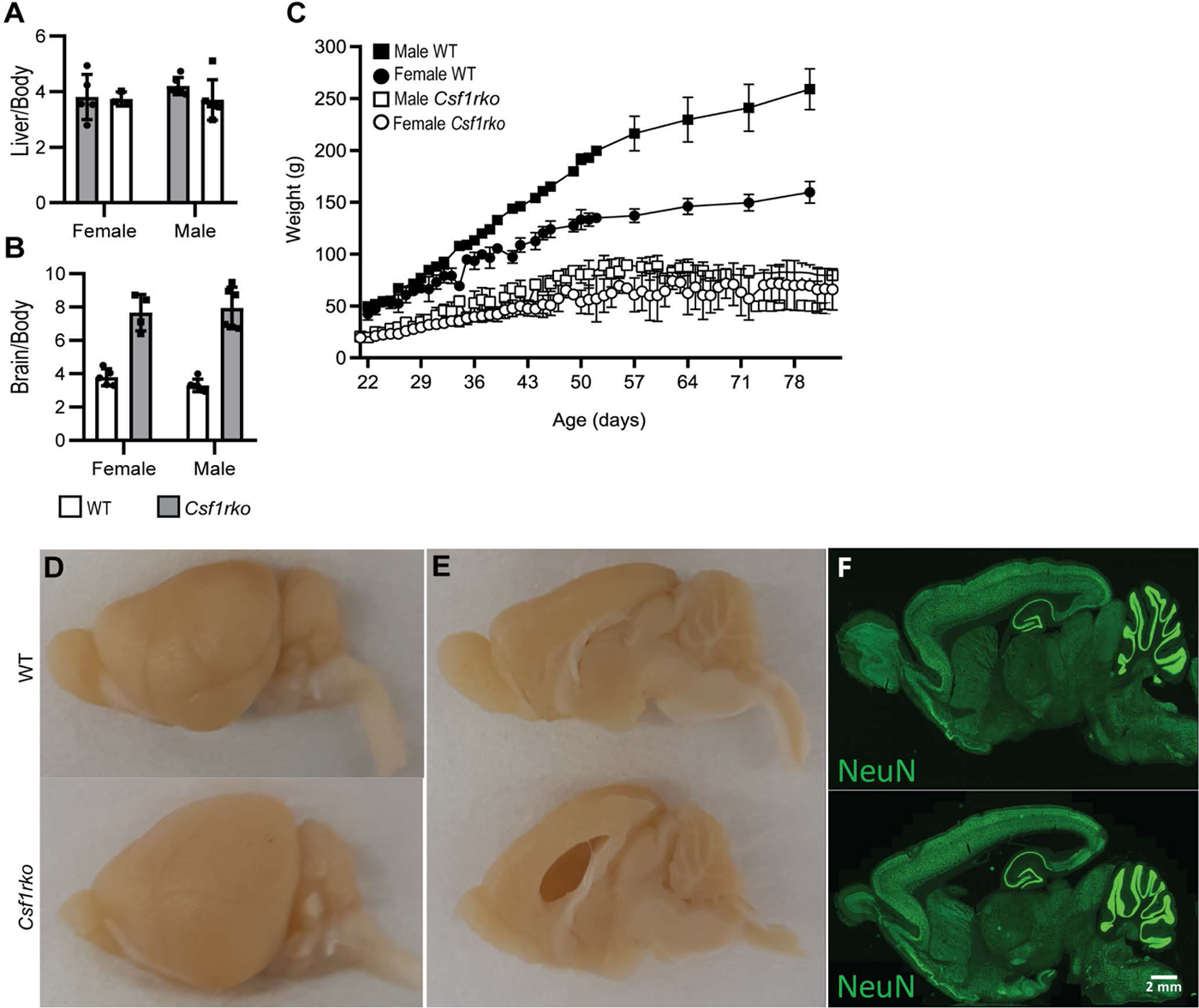
Liver, brain and body weight and brain morphology in *Csf1rko* and WT rats. (A) Liver/body and (B) brain/body weight ratio in 3 wk-old male and female WT and *Csf1rko* rats. (C) Time course of body weight gain of male and female WT and *Csf1rko* rats. (D) Representative images of WT (top panel) and *Csf1rko* (bottom panel) whole rat brain cut in the midsagittal plane. (E) Olfactory bulb, ventricular and cerebellar development of WT (top panel) and *Csf1rko* (bottom panel). (F) NeuN staining of sagittal slices from WT (top panel) and *Csf1rko* (bottom panel) brains including the cerebellum indicating neuronal distribution.

**Figure S2.**
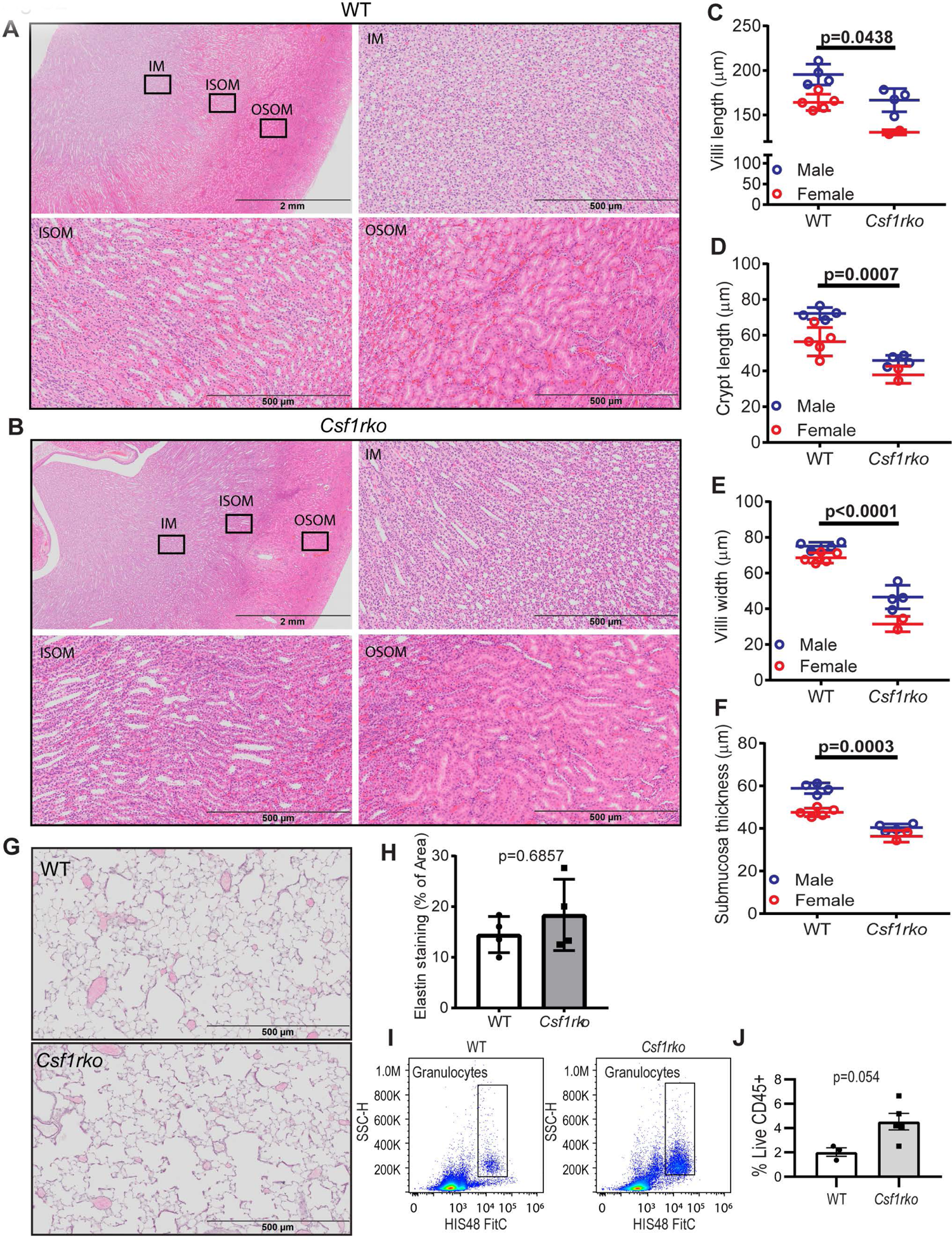
Histological analysis of *Csf1rko* and WT kidney, lung and intestine. (A, B) Representative images of H&E staining in the kidney of (A) WT and (B) *Csf1rko* rats. Inner medulla (IM) outer stripe of the outer medulla (OSOM) and inner stripe of the outer medulla (ISOM) are enlarged. (C-F) Quantitative morphometry of villus architecture of WT and *Csf1rko* rats performed at 3 wks of age as described in Materials and Methods. (G) Representative images of elastin staining in the lung of WT and *Csf1rko* rats. (H) quantitative analysis of elastin stained area (% of area) of the lung of WT and *Csf1rko* rats. (I) Flow cytometry gating strategy to identify granulocytes in disaggregated lung. (J) Quantitation of granulocytes in 9 wk-old WT and *Csf1rko* rat lungs. p value based upon Mann Whitney test.

**Figure S3.**
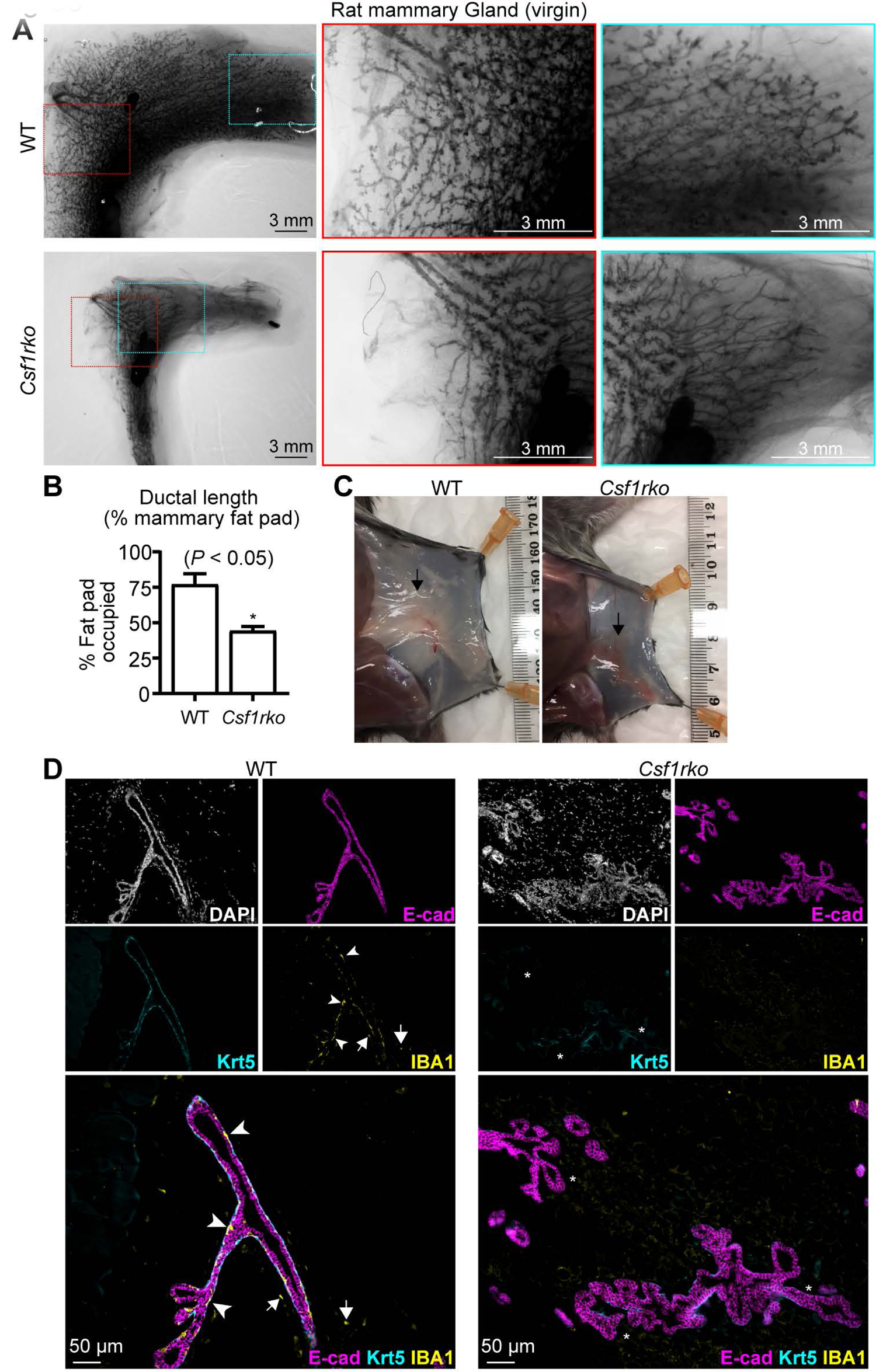
Comparison of mammary gland development in WT and *Csf1rko* rats. (A) Representative mammary gland wholemounts from 9.5-week-old WT and *Csf1rko* rats. (B) Ductal growth relative to growth of the mammary fat pad. * P < 0.05, Student’s t-test. (C) Photographs of rat mammary tissue in situ from WT and *Csf1rko* rats immediately post-euthanasia (black arrows). (D) IBA1 immunostaining of rat mammary tissue. E-cadherin (e-Cad) stains cells of the luminal lineage and cytokeratin 5 (Krt5) stains the basal lineage. Nuclei are stained with DAPI. White arrowheads point to intraepithelial IBA1^+^ macrophages; white arrows point to IBA1^+^ stromal macrophages; asterisks show regions in *Csf1rko* tissue with poor basal cell development. Representative of n = 3 rats from each group.

**Figure S4.**
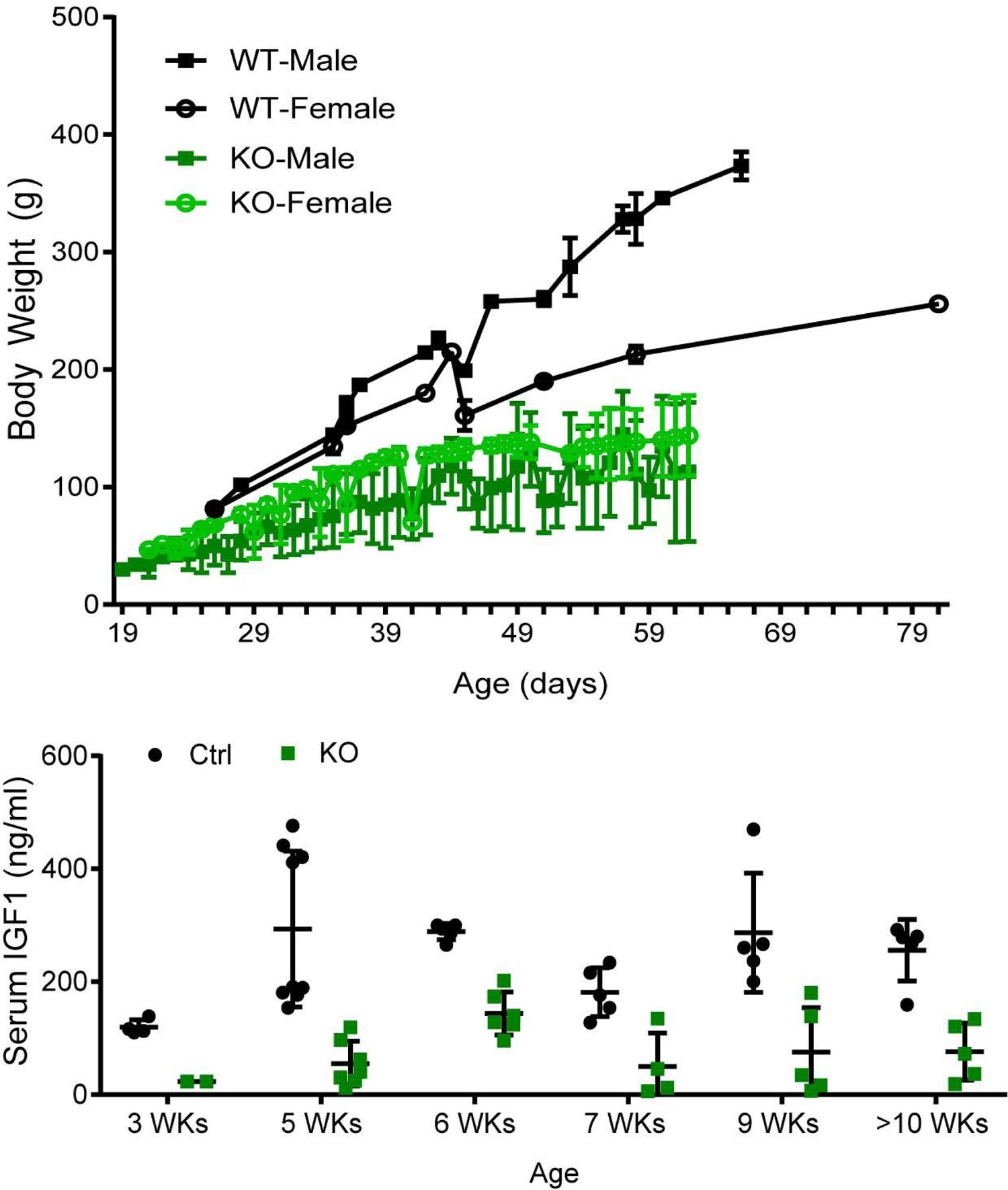
(A) Time course of postnatal body weight gain of outbred Sprague-Dawley (SD) male and female WT and *Csf1rko* rats. (B) Serum IGF-1 levels in a mixed cohort of male and female WT, *Csf1rko* SD rats.

**Figure S5.**
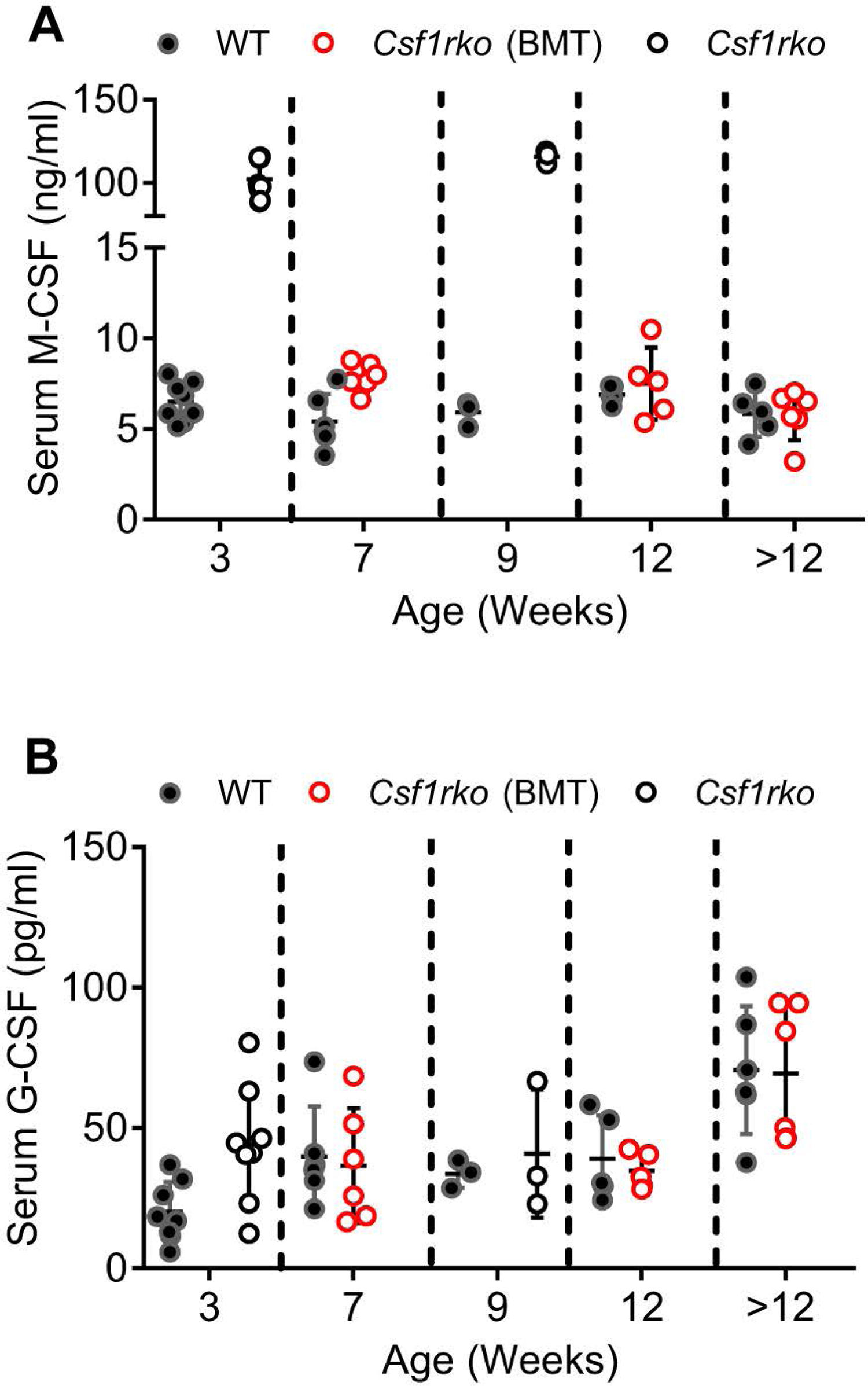
(A) Serum level of macrophage colony stimulating factor (CSF1) and (B) Granulocyte Colony Stimulating Factor (CSF3) in a mixed cohort of male and female WT, *Csf1rko* following bone marrow transfer (BMT) at 3 wks and *Csf1rko* rats at the ages indicated.

## Supplementary Table

Table S1. Comparative analysis of gene expression in the liver of WT and Csf1rko rats A. Primary output of Kallisto quantitation of gene expression for livers from 3 male and 3 female WT and *Csf1ro* rats at 3 wks. List is ranked based upon ratio of expression KO/WT. B. Analysis of differentially expressed genes (DEG) distinguishing WT and *Csf1rko* livers. C. Co-expressed gene clusters generated from Biolayout analysis. D. GO term enrichment for genes within the larger clusters of co-expressed genes. E. Expression profiles of growth-associated transcripts and cell-type specific markers in WT and *Csf1rko* livers

## References

1. Guilliams M, Thierry GR, Bonnardel J, Bajenoff M. Establishment and Maintenance of the Macrophage Niche. Immunity. 2020;52(3):434–51. Epub 2020/03/19. doi: 10.1016/j.immuni.2020.02.015. PubMed PMID: 32187515.

2. Hume DA, Irvine KM, Pridans C. The Mononuclear Phagocyte System: The Relationship between Monocytes and Macrophages. Trends Immunol. 2019;40:98–112. Epub 2018/12/24. doi: 10.1016/j.it.2018.11.007. PubMed PMID: 30579704.

3. Chitu V, Stanley ER. Regulation of Embryonic and Postnatal Development by the CSF-1 Receptor. Curr Top Dev Biol. 2017;123:229–75. Epub 2017/02/27. doi: 10.1016/bs.ctdb.2016.10.004. PubMed PMID: 28236968; PubMed Central PMCID: PMCPMC5479137.

4. Garceau V, Balic A, Garcia-Morales C, Sauter KA, McGrew MJ, Smith J, et al. The development and maintenance of the mononuclear phagocyte system of the chick is controlled by signals from the macrophage colony-stimulating factor receptor. BMC Biol. 2015;13:12. Epub 2015/04/11. doi: 10.1186/s12915-015-0121-9. PubMed PMID: 25857347; PubMed Central PMCID: PMCPMC4369834.

5. Garceau V, Smith J, Paton IR, Davey M, Fares MA, Sester DP, et al. Pivotal Advance: Avian colony-stimulating factor 1 (CSF-1), interleukin-34 (IL-34), and CSF-1 receptor genes and gene products. J Leukoc Biol. 2010;87(5):753–64. Epub 2010/01/07. doi: 10.1189/jlb.0909624. PubMed PMID: 20051473.

6. Hume DA, Caruso M, Ferrari-Cestari M, Summers KM, Pridans C, Irvine KM. Phenotypic impacts of CSF1R deficiencies in humans and model organisms. J Leukoc Biol. 2020;107(2):205–19. Epub 2019/07/23. doi: 10.1002/JLB.MR0519-143R. PubMed PMID: 31330095.

7. Wynn TA, Chawla A, Pollard JW. Macrophage biology in development, homeostasis and disease. Nature. 2013;496(7446):445-55. Epub 2013/04/27. doi: 10.1038/nature12034. PubMed PMID: 23619691; PubMed Central PMCID: PMCPMC3725458.

8. Guo L, Bertola DR, Takanohashi A, Saito A, Segawa Y, Yokota T, et al. Bi-allelic CSF1R Mutations Cause Skeletal Dysplasia of Dysosteosclerosis-Pyle Disease Spectrum and Degenerative Encephalopathy with Brain Malformation. Am J Hum Genet. 2019;104(5):925–35. Epub 2019/04/16. doi: 10.1016/j.ajhg.2019.03.004. PubMed PMID: 30982609.

9. Oosterhof N, Chang IJ, Karimiani EG, Kuil LE, Jensen DM, Daza R, et al. Homozygous Mutations in CSF1R Cause a Pediatric-Onset Leukoencephalopathy and Can Result in Congenital Absence of Microglia. Am J Hum Genet. 2019;104(5):936–47. Epub 2019/04/16. doi: 10.1016/j.ajhg.2019.03.010. PubMed PMID: 30982608; PubMed Central PMCID: PMCPMC6506793.

10. Kindis E, Simsek-Kiper PO, Kosukcu C, Taskiran EZ, Gocmen R, Utine E, et al. Further expanding the mutational spectrum of brain abnormalities, neurodegeneration, and dysosteosclerosis: A rare disorder with neurologic regression and skeletal features. Am J Med Genet A. 2021. Epub 2021/03/23. doi: 10.1002/ajmg.a.62179. PubMed PMID: 33749994.

11. Tamhankar PM, Zhu B, Tamhankar VP, Mithbawkar S, Seabra L, Livingston JH, et al. A Novel Hypomorphic CSF1R Gene Mutation in the Biallelic State Leading to Fatal Childhood Neurodegeneration. Neuropediatrics. 2020;51(4):302–6. Epub 2020/05/29. doi: 10.1055/s-0040-1702161. PubMed PMID: 32464672.

12. Hume DA, Caruso M, Keshvari S, Patkar OL, Sehgal A, Bush SJ, et al. The mononuclear phagocyte system of the rat. J Immunol. 2021:In press.

13. Pridans C, Raper A, David GM, Alves J, Sauter KA, Lefevre L, et al. Pleiotropic Impacts of Macrophage and Microglial Deficiency on Development in Rats with Targeted Mutation of the Csf1r Locus. J Immunol. 2018;201(9):2683–99. doi: 10.4049/jimmunol.1701783. Epub 2018 Sep 24. PMID: 30249809

14. Bennett FC, Bennett ML, Yaqoob F, Mulinyawe SB, Grant GA, Hayden Gephart M, et al. A Combination of Ontogeny and CNS Environment Establishes Microglial Identity. Neuron. 2018;98(6):1170–83 e8. Epub 2018/06/05. doi: 10.1016/j.neuron.2018.05.014. PubMed PMID: 29861285; PubMed Central PMCID: PMCPMC6023731.

15. Erblich B, Zhu L, Etgen AM, Dobrenis K, Pollard JW. Absence of colony stimulation factor-1 receptor results in loss of microglia, disrupted brain development and olfactory deficits. PLoS One. 2011;6(10):e26317. Epub 2011/11/03. doi: 10.1371/journal.pone.0026317. PubMed PMID: 22046273; PubMed Central PMCID: PMCPMC3203114.

16. Henson PM, Hume DA. Apoptotic cell removal in development and tissue homeostasis. Trends Immunol. 2006;27(5):244–50. Epub 2006/04/06. doi: 10.1016/j.it.2006.03.005. PubMed PMID: 16584921.

17. Lichanska AM, Browne CM, Henkel GW, Murphy KM, Ostrowski MC, McKercher SR, et al. Differentiation of the mononuclear phagocyte system during mouse embryogenesis: the role of transcription factor PU.1. Blood. 1999;94(1):127-38. Epub 1999/06/25. PubMed PMID: 10381505.

18. Patkar OL, Caruso M, Teakle N, Keshvari S, Bush SJ, Pridans C, et al. Analysis of homozygous and heterozygous Csf1r knockout in the rat as a model for understanding microglial function in brain development and the impacts of human CSF1R mutations. Neurobiol Dis. 2021:105268. Epub 2021/01/16. doi: 10.1016/j.nbd.2021.105268. PubMed PMID: 33450391.

19. Oosterhof N, Kuil LE, van der Linde HC, Burm SM, Berdowski W, van Ijcken WFJ, et al. Colony-Stimulating Factor 1 Receptor (CSF1R) Regulates Microglia Density and Distribution, but Not Microglia Differentiation In Vivo. Cell Rep. 2018;24(5):1203–17 e6. Epub 2018/08/02. doi: 10.1016/j.celrep.2018.06.113. PubMed PMID: 30067976.

20. Michalopoulos GK. Hepatostat: Liver regeneration and normal liver tissue maintenance. Hepatology. 2017;65(4):1384–92. Epub 2016/12/21. doi: 10.1002/hep.28988. PubMed PMID: 27997988.

21. Michalopoulos GK, Bhushan B. Liver regeneration: biological and pathological mechanisms and implications. Nat Rev Gastroenterol Hepatol. 2021;18(1):40–55. Epub 2020/08/09. doi: 10.1038/s41575-020-0342-4. PubMed PMID: 32764740.

22. Pridans C, Irvine KM, Davis GM, Lefevre L, Bush SJ, Hume DA. Transcriptomic Analysis of Rat Macrophages. Front Immunol. 2020;11:594594. Epub 2021/02/27. doi: 10.3389/fimmu.2020.594594. PubMed PMID: 33633725; PubMed Central PMCID: PMCPMC7902030.

23. Alexander B, Guzail MA, Foster CS. Morphological changes during hepatocellular maturity in neonatal rats. Anat Rec. 1997;248(1):104–9. Epub 1997/05/01. doi: 10.1002/(SICI)1097-0185(199705)248:1&104::AID-AR12>3.0.CO;2-T. PubMed PMID: 9143673.

24. Septer S, Edwards G, Gunewardena S, Wolfe A, Li H, Daniel J, et al. Yes-associated protein is involved in proliferation and differentiation during postnatal liver development. Am J Physiol Gastrointest Liver Physiol. 2012;302(5):G493-503. Epub 2011/12/24. doi: 10.1152/ajpgi.00056.2011. PubMed PMID: 22194415; PubMed Central PMCID: PMCPMC3311431.

25. Summers KM, Hume DA. Identification of the macrophage-specific promoter signature in FANTOM5 mouse embryo developmental time course data. J Leukoc Biol. 2017;102(4):1081–92. Epub 2017/07/29. doi: 10.1189/jlb.1A0417-150RR. PubMed PMID: 28751473.

26. Lichanska AM, Waters MJ. How growth hormone controls growth, obesity and sexual dimorphism. Trends Genet. 2008;24(1):41–7. Epub 2007/12/08. doi: 10.1016/j.tig.2007.10.006. PubMed PMID: 18063438.

27. Pridans C, Sauter KA, Irvine KM, Davis GM, Lefevre L, Raper A, et al. Macrophage colony-stimulating factor increases hepatic macrophage content, liver growth, and lipid accumulation in neonatal rats. Am J Physiol Gastrointest Liver Physiol. 2018;314(3):G388-G98. Epub 2018/01/20. doi: 10.1152/ajpgi.00343.2017. PubMed PMID: 29351395; PubMed Central PMCID: PMCPMC5899243.

28. Banaei-Bouchareb L, Gouon-Evans V, Samara-Boustani D, Castellotti MC, Czernichow P, Pollard JW, et al. Insulin cell mass is altered in Csf1op/Csf1op macrophage-deficient mice. J Leukoc Biol. 2004;76(2):359–67. Epub 2004/06/05. doi: 10.1189/jlb.1103591. PubMed PMID: 15178709.

29. Albertoni Borghese MF, Ortiz MC, Balonga S, Moreira Szokalo R, Majowicz MP. The Role of Endothelin System in Renal Structure and Function during the Postnatal Development of the Rat Kidney. PLoS One. 2016;11(2):e0148866. Epub 2016/02/13. doi: 10.1371/journal.pone.0148866. PubMed PMID: 26872270; PubMed Central PMCID: PMCPMC4752218.

30. Sehgal A, Donaldson DS, Pridans C, Sauter KA, Hume DA, Mabbott NA. The role of CSF1R-dependent macrophages in control of the intestinal stem-cell niche. Nat Commun. 2018;9(1):1272. Epub 2018/03/30. doi: 10.1038/s41467-018-03638-6. PubMed PMID: 29593242; PubMed Central PMCID: PMCPMC5871851.

31. Stewart TA, Hughes K, Hume DA, Davis FM. Developmental Stage-Specific Distribution of Macrophages in Mouse Mammary Gland. Front Cell Dev Biol. 2019;7:250. Epub 2019/11/12. doi: 10.3389/fcell.2019.00250. PubMed PMID: 31709255; PubMed Central PMCID: PMCPMC6821639.

32. Irvine KM, Caruso M, Cestari MF, Davis GM, Keshvari S, Sehgal A, et al. Analysis of the impact of CSF-1 administration in adult rats using a novel Csf1r-mApple reporter gene. J Leukoc Biol. 2020;107(2):221–35. Epub 2019/08/10. doi: 10.1002/JLB.MA0519-149R. PubMed PMID: 31397014.

33. Summers KM, Bush SJ, Hume DA. Transcriptional network analysis of transcriptomic diversity in resident tissue macrophages and dendritic cells in the mouse mononuclear phagocyte system. PLoS Biol. 2020 Oct 8;18(10):e3000859. doi: 10.1371/journal.pbio.3000859. eCollection 2020 Oct. PMID: 33031383

34. Gow DJ, Sester DP, Hume DA. CSF-1, IGF-1, and the control of postnatal growth and development. J Leukoc Biol. 2010;88(3):475-81. Epub 2010/06/04. doi: 10.1189/jlb.0310158. PubMed PMID: 20519640.

35. Kohler C. Allograft inflammatory factor-1/Ionized calcium-binding adapter molecule 1 is specifically expressed by most subpopulations of macrophages and spermatids in testis. Cell Tissue Res. 2007;330(2):291–302. Epub 2007/09/18. doi: 10.1007/s00441-007-0474-7. PubMed PMID: 17874251.

36. Rojo R, Raper A, Ozdemir DD, Lefevre L, Grabert K, Wollscheid-Lengeling E, et al. Deletion of a Csf1r enhancer selectively impacts CSF1R expression and development of tissue macrophage populations. Nat Commun. 2019;10(1):3215. Epub 2019/07/22. doi: 10.1038/s41467-019-11053-8. PubMed PMID: 31324781; PubMed Central PMCID: PMCPMC6642117.

37. Dai XM, Ryan GR, Hapel AJ, Dominguez MG, Russell RG, Kapp S, et al. Targeted disruption of the mouse colony-stimulating factor 1 receptor gene results in osteopetrosis, mononuclear phagocyte deficiency, increased primitive progenitor cell frequencies, and reproductive defects. Blood. 2002;99(1):111–20. Epub 2002/01/05. doi: 10.1182/blood.v99.1.111. PubMed PMID: 11756160.

38. Kaur S, Raggatt LJ, Batoon L, Hume DA, Levesque JP, Pettit AR. Role of bone marrow macrophages in controlling homeostasis and repair in bone and bone marrow niches. Semin Cell Dev Biol. 2017;61:12–21. Epub 2016/08/16. doi: 10.1016/j.semcdb.2016.08.009. PubMed PMID: 27521519.

39. Bush SJ, McCulloch MEB, Lisowski ZM, Muriuki C, Clark EL, Young R, et al. Species-Specificity of Transcriptional Regulation and the Response to Lipopolysaccharide in Mammalian Macrophages. Front Cell Dev Biol. 2020;8:661. Epub 2020/08/15. doi: 10.3389/fcell.2020.00661. PubMed PMID: 32793601; PubMed Central PMCID: PMCPMC7386301.

40. Baker J, Liu JP, Robertson EJ, Efstratiadis A. Role of insulin-like growth factors in embryonic and postnatal growth. Cell. 1993;75(1):73–82. Epub 1993/10/08. PubMed PMID: 8402902.

41. Monies D, Maddirevula S, Kurdi W, Alanazy MH, Alkhalidi H, Al-Owain M, et al. Autozygosity reveals recessive mutations and novel mechanisms in dominant genes: implications in variant interpretation. Genet Med. 2017;19(10):1144–50. Epub 2017/04/07. doi: 10.1038/gim.2017.22. PubMed PMID: 28383543.

42. Kineman RD, Del Rio-Moreno M, Sarmento-Cabral A. 40 YEARS of IGF1: Understanding the tissue-specific roles of IGF1/IGF1R in regulating metabolism using the Cre/loxP system. J Mol Endocrinol. 2018;61(1):T187–T98. Epub 2018/05/11. doi: 10.1530/JME-18-0076. PubMed PMID: 29743295.

43. Duran-Ortiz S, Noboa V, Kopchick JJ. Tissue-specific disruption of the growth hormone receptor (GHR) in mice: An update. Growth Horm IGF Res. 2020;51:1–5. Epub 2020/01/11. doi: 10.1016/j.ghir.2019.12.004. PubMed PMID: 31923746.

44. Bikle DD, Tahimic C, Chang W, Wang Y, Philippou A, Barton ER. Role of IGF-I signaling in muscle bone interactions. Bone. 2015;80:79–88. Epub 2015/10/11. doi: 10.1016/j.bone.2015.04.036. PubMed PMID: 26453498; PubMed Central PMCID: PMCPMC4600536.

45. Davies JS, Gevers EF, Stevenson AE, Coschigano KT, El-Kasti MM, Bull MJ, et al. Adiposity profile in the dwarf rat: an unusually lean model of profound growth hormone deficiency. Am J Physiol Endocrinol Metab. 2007;292(5):E1483–94. Epub 2007/02/01. doi: 10.1152/ajpendo.00417.2006. PubMed PMID: 17264226.

46. Coschigano KT, Holland AN, Riders ME, List EO, Flyvbjerg A, Kopchick JJ. Deletion, but not antagonism, of the mouse growth hormone receptor results in severely decreased body weights, insulin, and insulin-like growth factor I levels and increased life span. Endocrinology. 2003;144(9):3799–810. Epub 2003/08/23. doi: 10.1210/en.2003-0374. PubMed PMID: 12933651.

47. Aguiar-Oliveira MH, Bartke A. Growth Hormone Deficiency: Health and Longevity. Endocr Rev. 2019;40(2):575-601. Epub 2018/12/24. doi: 10.1210/er.2018-00216. PubMed PMID: 30576428; PubMed Central PMCID: PMCPMC6416709.

48. Vannella KM, Wynn TA. Mechanisms of Organ Injury and Repair by Macrophages. Annu Rev Physiol. 2017;79:593–617. Epub 2016/12/14. doi: 10.1146/annurev-physiol-022516-034356. PubMed PMID: 27959618.

49. Chang HR, Kim HJ, Xu X, Ferrante AW, Jr. Macrophage and adipocyte IGF1 maintain adipose tissue homeostasis during metabolic stresses. Obesity (Silver Spring). 2016;24(1):172–83. Epub 2015/12/15. doi: 10.1002/oby.21354. PubMed PMID: 26663512; PubMed Central PMCID: PMCPMC4793714.

50. Cotton WR, Gaines JF. Unerupted dentition secondary to congenital osteopetrosis in the Osborne-Mendel rat. Proc Soc Exp Biol Med. 1974;146(2):554–61. Epub 1974/06/01. doi: 10.3181/00379727-146-38146. PubMed PMID: 4834464.

51. Van Wesenbeeck L, Odgren PR, MacKay CA, D’Angelo M, Safadi FF, Popoff SN, et al. The osteopetrotic mutation toothless (tl) is a loss-of-function frameshift mutation in the rat Csf1 gene: Evidence of a crucial role for CSF-1 in osteoclastogenesis and endochondral ossification. Proc Natl Acad Sci U S A. 2002;99(22):14303–8. Epub 2002/10/16. doi: 10.1073/pnas.202332999. PubMed PMID: 12379742; PubMed Central PMCID: PMCPMC137879.

52. Mass E, Ballesteros I, Farlik M, Halbritter F, Gunther P, Crozet L, et al. Specification of tissue-resident macrophages during organogenesis. Science. 2016;353(6304). Epub 2016/08/06. doi: 10.1126/science.aaf4238. PubMed PMID: 27492475; PubMed Central PMCID: PMCPMC5066309.

53. Scott CL, Zheng F, De Baetselier P, Martens L, Saeys Y, De Prijck S, et al. Bone marrow-derived monocytes give rise to self-renewing and fully differentiated Kupffer cells. Nat Commun. 2016;7:10321. Epub 2016/01/28. doi: 10.1038/ncomms10321. PubMed PMID: 26813785; PubMed Central PMCID: PMCPMC4737801.

54. Gow DJ, Sauter KA, Pridans C, Moffat L, Sehgal A, Stutchfield BM, et al. Characterisation of a novel Fc conjugate of macrophage colony-stimulating factor. Mol Ther. 2014;22(9):1580–92. Epub 2014/06/26. doi: 10.1038/mt.2014.112. PubMed PMID: 24962162; PubMed Central PMCID: PMCPMC4435485.

55. Sauter KA, Waddell LA, Lisowski ZM, Young R, Lefevre L, Davis GM, et al. Macrophage colony-stimulating factor (CSF1) controls monocyte production and maturation and the steady-state size of the liver in pigs. Am J Physiol Gastrointest Liver Physiol. 2016;311(3):G533-47. Epub 2016/07/23. doi: 10.1152/ajpgi.00116.2016. PubMed PMID: 27445344; PubMed Central PMCID: PMCPMC5076001.

56. Stutchfield BM, Antoine DJ, Mackinnon AC, Gow DJ, Bain CC, Hawley CA, et al. CSF1 Restores Innate Immunity After Liver Injury in Mice and Serum Levels Indicate Outcomes of Patients With Acute Liver Failure. Gastroenterology. 2015;149(7):1896–909 e14. Epub 2015/09/08. doi: 10.1053/j.gastro.2015.08.053. PubMed PMID: 26344055; PubMed Central PMCID: PMCPMC4672154.

57. Peinado-Onsurbe J, Staels B, Deeb S, Ramirez I, Llobera M, Auwerx J. Neonatal extinction of liver lipoprotein lipase expression. Biochim Biophys Acta. 1992;1131(3):281–6. Epub 1992/07/15. doi: 10.1016/0167-4781(92)90026-v. PubMed PMID: 1627643.

58. Kammerer L, Mohammad GH, Wolna M, Robbins P, Lakhal-Littleton S. Fetal liver hepcidin secures iron stores in utero. Blood. 2020. Epub 2020/06/17. doi: 10.1182/blood.2019003907. PubMed PMID: 32542311.

59. Cramer S, Beveridge M, Kilberg M, Novak D. Physiological importance of system A-mediated amino acid transport to rat fetal development. Am J Physiol Cell Physiol. 2002;282(1):C153–60. Epub 2001/12/18. doi: 10.1152/ajpcell.2002.282.1.C153. PubMed PMID: 11742808.

60. Fuhrmeister J, Zota A, Sijmonsma TP, Seibert O, Cingir S, Schmidt K, et al. Fasting-induced liver GADD45beta restrains hepatic fatty acid uptake and improves metabolic health. EMBO Mol Med. 2016;8(6):654–69. Epub 2016/05/04. doi: 10.15252/emmm.201505801. PubMed PMID: 27137487; PubMed Central PMCID: PMCPMC4888855.

61. Shi H, Yao R, Lian S, Liu P, Liu Y, Yang YY, et al. Regulating glycolysis, the TLR4 signal pathway and expression of RBM3 in mouse liver in response to acute cold exposure. Stress. 2019;22(3):366–76. Epub 2019/03/02. doi: 10.1080/10253890.2019.1568987. PubMed PMID: 30821572.

62. Adams JM, Otero-Corchon V, Hammond GL, Veldhuis JD, Qi N, Low MJ. Somatostatin is essential for the sexual dimorphism of GH secretion, corticosteroid-binding globulin production, and corticosterone levels in mice. Endocrinology. 2015;156(3):1052–65. Epub 2015/01/01. doi: 10.1210/en.2014-1429. PubMed PMID: 25551181; PubMed Central PMCID: PMCPMC4330306.

63. Martinez-Jimenez CP, Kyrmizi I, Cardot P, Gonzalez FJ, Talianidis I. Hepatocyte nuclear factor 4alpha coordinates a transcription factor network regulating hepatic fatty acid metabolism. Mol Cell Biol. 2010;30(3):565–77. Epub 2009/11/26. doi: 10.1128/MCB.00927-09. PubMed PMID: 19933841; PubMed Central PMCID: PMCPMC2812226.

64. Apte U, Zeng G, Thompson MD, Muller P, Micsenyi A, Cieply B, et al. beta-Catenin is critical for early postnatal liver growth. Am J Physiol Gastrointest Liver Physiol. 2007;292(6):G1578–85. Epub 2007/03/03. doi: 10.1152/ajpgi.00359.2006. PubMed PMID: 17332475.

65. Lupu F, Terwilliger JD, Lee K, Segre GV, Efstratiadis A. Roles of growth hormone and insulin-like growth factor 1 in mouse postnatal growth. Dev Biol. 2001;229(1):141–62. Epub 2001/01/03. doi: 10.1006/dbio.2000.9975. PubMed PMID: 11133160.

66. Percival CJ, Richtsmeier JT. Angiogenesis and intramembranous osteogenesis. Dev Dyn. 2013;242(8):909-22. Epub 2013/06/06. doi: 10.1002/dvdy.23992. PubMed PMID: 23737393; PubMed Central PMCID: PMCPMC3803110.

67. Opperman LA. Cranial sutures as intramembranous bone growth sites. Dev Dyn. 2000;219(4):472–85. Epub 2000/11/21. doi: 10.1002/1097-0177(2000) PubMed PMID: 11084647.

68. Chang MK, Raggatt LJ, Alexander KA, Kuliwaba JS, Fazzalari NL, Schroder K, et al. Osteal tissue macrophages are intercalated throughout human and mouse bone lining tissues and regulate osteoblast function in vitro and in vivo. J Immunol. 2008;181(2):1232–44. Epub 2008/07/09. doi: 10.4049/jimmunol.181.2.1232. PubMed PMID: 18606677.

69. Alexander KA, Chang MK, Maylin ER, Kohler T, Muller R, Wu AC, et al. Osteal macrophages promote in vivo intramembranous bone healing in a mouse tibial injury model. J Bone Miner Res. 2011;26(7):1517–32. Epub 2011/02/10. doi: 10.1002/jbmr.354. PubMed PMID: 21305607.

70. Batoon L, Millard SM, Wullschleger ME, Preda C, Wu AC, Kaur S, et al. CD169(+) macrophages are critical for osteoblast maintenance and promote intramembranous and endochondral ossification during bone repair. Biomaterials. 2019;196:51–66. Epub 2017/11/07. doi: 10.1016/j.biomaterials.2017.10.033. PubMed PMID: 29107337.

71. Watanabe H, MacKay CA, Mason-Savas A, Kislauskis EH, Marks SC, Jr. Colony-stimulating factor-1 (CSF-1) rescues osteoblast attachment, survival and sorting of beta-actin mRNA in the toothless (tl-osteopetrotic) mutation in the rat. Int J Dev Biol. 2000;44(2):201–7. Epub 2000/05/04. PubMed PMID: 10794078.

72. Lazarus S, Tseng HW, Lawrence F, Woodruff MA, Duncan EL, Pettit AR. Characterization of Normal Murine Carpal Bone Development Prompts Re-Evaluation of Pathologic Osteolysis as the Cause of Human Carpal-Tarsal Osteolysis Disorders. Am J Pathol. 2017;187(9):1923–34. Epub 2017/07/05. doi: 10.1016/j.ajpath.2017.05.007. PubMed PMID: 28675805.

73. Sobacchi C, Schulz A, Coxon FP, Villa A, Helfrich MH. Osteopetrosis: genetics, treatment and new insights into osteoclast function. Nat Rev Endocrinol. 2013;9(9):522–36. Epub 2013/07/24. doi: 10.1038/nrendo.2013.137. PubMed PMID: 23877423.

74. MacDonald KP, Palmer JS, Cronau S, Seppanen E, Olver S, Raffelt NC, et al. An antibody against the colony-stimulating factor 1 receptor depletes the resident subset of monocytes and tissue- and tumor-associated macrophages but does not inhibit inflammation. Blood. 2010;116(19):3955–63. Epub 2010/08/05. doi: 10.1182/blood-2010-02-266296. PubMed PMID: 20682855.

75. Kaur S, Raggatt LJ, Millard SM, Wu AC, Batoon L, Jacobsen RN, et al. Self-repopulating recipient bone marrow resident macrophages promote long-term hematopoietic stem cell engraftment. Blood. 2018;132(7):735–49. Epub 2018/06/28. doi: 10.1182/blood-2018-01-829663. PubMed PMID: 29945953.

76. Norgard M, Marks SC, Jr., Reinholt FP, Andersson G. The effects of colony-stimulating factor-1 (CSF-1) on the development of osteoclasts and their expression of tartrate-resistant acid phosphatase (TRAP) in toothless (tl-osteopetrotic) rats. Crit Rev Eukaryot Gene Expr. 2003;13(2-4):117–32. Epub 2003/12/31. doi: 10.1615/critreveukaryotgeneexpr.v13.i24.60. PubMed PMID: 14696961.

77. Hidalgo A, Chilvers ER, Summers C, Koenderman L. The Neutrophil Life Cycle. Trends Immunol. 2019;40(7):584–97. Epub 2019/06/04. doi: 10.1016/j.it.2019.04.013. PubMed PMID: 31153737.

78. Shi J, Gilbert GE, Kokubo Y, Ohashi T. Role of the liver in regulating numbers of circulating neutrophils. Blood. 2001;98(4):1226–30. Epub 2001/08/09. doi: 10.1182/blood.v98.4.1226. PubMed PMID: 11493474.

79. Massberg S, Schaerli P, Knezevic-Maramica I, Kollnberger M, Tubo N, Moseman EA, et al. Immunosurveillance by hematopoietic progenitor cells trafficking through blood, lymph, and peripheral tissues. Cell. 2007;131(5):994–1008. Epub 2007/11/30. doi: 10.1016/j.cell.2007.09.047. PubMed PMID: 18045540; PubMed Central PMCID: PMCPMC2330270.

80. Hettinger J, Richards DM, Hansson J, Barra MM, Joschko AC, Krijgsveld J, et al. Origin of monocytes and macrophages in a committed progenitor. Nat Immunol. 2013;14(8):821–30. Epub 2013/07/03. doi: 10.1038/ni.2638. PubMed PMID: 23812096.

81. Yahara Y, Barrientos T, Tang YJ, Puviindran V, Nadesan P, et al. Erythromyeloid progenitors give rise to a population of osteoclasts that contribute to bone homeostasis and repair. Nat Cell Biol. 2020 Jan;22(1):49–59. doi: 10.1038/s41556-019-0437-8. Epub 2020 Jan 6. PMID: 31907410

82. Grabert K, Sehgal A, Irvine KM, Wollscheid-Lengeling E, Ozdemir DD, Stables J, et al. A mouse transgenic line that reports CSF1R protein expression provides a definitive differentiation marker for the mouse mononuclear phagocyte system. J Immunol. 2020 Dec 1;205(11):3154–3166. doi: 10.4049/jimmunol.2000835. Epub 2020 Nov 2. PMID: 33139489

83. Wu J, Wang C, Li S, Li S, Wang W, Li J, et al. Thyroid hormone-responsive SPOT 14 homolog promotes hepatic lipogenesis, and its expression is regulated by liver X receptor alpha through a sterol regulatory element-binding protein 1c-dependent mechanism in mice. Hepatology. 2013;58(2):617–28. Epub 2013/01/26. doi: 10.1002/hep.26272. PubMed PMID: 23348573.

84. Wallin J, Wilting J, Koseki H, Fritsch R, Christ B, Balling R. The role of Pax-1 in axial skeleton development. Development. 1994;120(5):1109–21. Epub 1994/05/01. PubMed PMID: 8026324.

85. Chen S, Zhou Y, Chen Y, Gu J. fastp: an ultra-fast all-in-one FASTQ preprocessor. Bioinformatics. 2018;34(17):i884–i90. Epub 2018/11/14. doi: 10.1093/bioinformatics/bty560. PubMed PMID: 30423086; PubMed Central PMCID: PMCPMC6129281.

86. Bray NL, Pimentel H, Melsted P, Pachter L. Near-optimal probabilistic RNA-seq quantification. Nat Biotechnol. 2016;34(5):525–7. Epub 2016/04/05. doi: 10.1038/nbt.3519. PubMed PMID: 27043002.

87. Steyn FJ, Huang L, Ngo ST, Leong JW, Tan HY, Xie TY, et al. Development of a method for the determination of pulsatile growth hormone secretion in mice. Endocrinology. 2011;152(8):3165–71. Epub 2011/05/19. doi: 10.1210/en.2011-0253. PubMed PMID: 21586549.

88. Sauter KA, Pridans C, Sehgal A, Tsai YT, Bradford BM, Raza S, et al. Pleiotropic effects of extended blockade of CSF1R signaling in adult mice. J Leukoc Biol. 2014;96(2):265–74. Epub 2014/03/22. doi: 10.1189/jlb.2A0114-006R. PubMed PMID: 24652541; PubMed Central PMCID: PMCPMC4378363.

89. Lloyd-Lewis B, Davis FM, Harris OB, Hitchcock JR, Lourenco FC, Pasche M, et al. Imaging the mammary gland and mammary tumours in 3D: optical tissue clearing and immunofluorescence methods. Breast Cancer Res. 2016;18(1):127. Epub 2016/12/15. doi: 10.1186/s13058-016-0754-9. PubMed PMID: 27964754; PubMed Central PMCID: PMCPMC5155399.

90. Davis FM, Lloyd-Lewis B, Harris OB, Kozar S, Winton DJ, Muresan L, et al. Single-cell lineage tracing in the mammary gland reveals stochastic clonal dispersion of stem/progenitor cell progeny. Nat Commun. 2016;7:13053. Epub 2016/10/26. doi: 10.1038/ncomms13053. PubMed PMID: 27779190; PubMed Central PMCID: PMCPMC5093309.

